# Assembly-free and alignment-free sample identification using genome skims

**DOI:** 10.1101/230409

**Authors:** Shahab Sarmashghi, Kristine Bohmann, M. Thomas P. Gilbert, Vineet Bafna, Siavash Mirarab

**Affiliations:** Department of Electrical & Computer Engineering, University of California, San Diego, La Jolla, CA 92093, USA; Evolutionary Genomics, Natural History Museum of Denmark, University of Copenhagen, Copenhagen, Denmark; School of Biological Sciences, University of East Anglia, Norwich, Norfolk, UK; Norwegian University of Science and Technology, University Museum, 7491 Trondheim, Norway; Department of Computer Science & Engineering, University of California, San Diego, La Jolla, CA 92093, USA

**Keywords:** Assembly-free, Alignment-free, DNA Barcoding, Genome-skimming, DNA reference databases, Next generation sequencing

## Abstract

The ability to quickly and inexpensively describe taxonomic diversity is critical in this era of rapid climate and biodiversity changes. The currently preferred molecular technique, barcoding, has been very successful, but is based on short organelle markers. Recently, an alternative *genome-skimming* approach has been proposed: low-pass sequencing (100Mb – several Gb per sample) is applied to voucher and/or query samples, and marker genes and/or organelle genomes are recovered computationally. The current practice of genome-skimming discards the vast majority of the data because the low coverage of genome-skims prevents assembling the nuclear genomes. In contrast, we suggest using all unassembled reads directly, but existing methods poorly support this goal. We introduce a new alignment-free tool, Skmer, to estimate genomic distances between the query and each reference genome-skim using the k-mer decomposition of reads. We test Skmer on a large set of insect and bird genomes, sub-sampled to create genome-skims. Skmer shows great accuracy in estimating genomic distances, identifying the closest match in a reference dataset, and inferring the phylogeny. The software is publicly available on https://github.com/shahab-sarmashghi/Skmer.git

## Background

The ability to quickly and inexpensively study the taxonomic diversity in an environment is critical in this era of rapid climate and biodiversity changes. The current molecular technique of choice is (meta)barcoding [1–3]. Traditional (meta)barcoding is based on DNA sequencing of taxonomically informative and group-specific marker genes (e.g., mitochondrial COI [1, 4] and 12S/16S [5, 6] for animals, chloroplast genes like matK for plants [7], and ITS [8] for fungi) that are variable enough for taxonomic identification, but have flanking regions that are sufficiently conserved to allow for PCR amplification using universal primers. Barcoding is used for taxonomic identification of single-species samples. In the case of metabarcoding, the goal is to deconstruct the taxonomic composition of a mixed sample consisting of multiple species [3]. Beyond the barcoding application, the barcoding marker genes have also been used to delimitate species [9] and to infer phylogenies [10, 11].

The accuracy of (meta)barcoding depends on the coverage of the reference database and the method used to search queries against it [3]. To increase coverage, reference databases with millions of barcodes have been generated (e.g., Barcode of Life Data System, BOLD, for COI [12]). Computational methods for finding the closest match in a reference dataset (e.g., TaxI [13]), and for placement of a query into existing marker trees [14–16] have been developed. However, the traditional approach to (meta)barcoding, despite its success, has some drawbacks. PCR for marker gene amplification requires relatively high quality DNA and thus cannot be applied to samples in which the DNA is heavily fragmented. Moreover, since barcode markers are relatively short regions, their phylogenetic signal and identification resolution can be limited [17]. For example, in a recent study, 896 out of 4,174 wasp species could not be distinguished from each other using COI barcodes [18].

While low costs have kept PCR-based pipelines attractive, decreasing costs of shotgun sequencing have now made it possible to shotgun sequence 1-2Gb of total DNA per reference specimen sample for as low as $80 [19], even after including sample preparation and labor costs. This has lead researchers to propose an alternate method that uses low-pass sequencing to generate *genome-skims* [19, 20], and subsequently identifies chloroplast or mitochondrial marker genes or assembles the organelle genome. Reconstructing plastid and mtDNA genomes from low-pass shotgun data is possible because organelle DNA tends to be heavily overrepresented in shotgun sequencing data; for example, 10.4% of all reads from the Apocynaceae family of flowering plants were from the chloroplast in one genome-skimming study [20]. Large reference databases based on genome-skimming techniques are under construction by projects such as PhyloAlps [21], NorBol [22], and DNAmark [23].

Most current applications of genome-skimming to species identification require organelle genome assembly, a task that requires relatively time-consuming manual curation steps to ensure that assembly errors are avoided [24]. This approach discards a vast proportion of the non-target data, reducing the discriminatory power. For these reasons, the DNAmark project [23] is considering alternative methods, where, instead of only relying on organelle markers, one could use the entire set of reads generated in a genome-skim as the identifier of a species. This approach poses an interesting methodological question: can the unassembled data be used to taxonomically profile reference and query samples in a similar manner to conventional barcoding, but using all available genomic information and saving us from the labor-intensive task of mitochondria/plastid genome assembly? In this paper, we introduce a new assembly-free method to directly use low coverage genome-skims of both reference and query samples. By avoiding the assembly step, our approach also reduces the amount of data processing needed for expanding the reference database.

We treat genome-skims simply as low-coverage “ bags of reads”, both for a collection of reference species and for query samples. The problem is to find the reference genome-skim that matches the query; if an exact match is not found, we seek the closest available match. A more advanced problem, not directly addressed here, is placing the query in a phylogeny of reference species. An even more difficult challenge, also not addressed here, is decomposing a query genome-skim that contains DNA from several different taxa into its constituent species.

Central to solving these problems is the ability to estimate a *distance* between two genome-skims for low and varied coverage using assembly-free and alignment-free approaches. Alignment-free sequence comparison has been widely studied [25–30], including for phylogenetic reconstruction [25, 31–43]. Most existing methods, such as Kr [28], andi [41], kmacs [44], and FSWM [43], compute evolutionary distances using the length distribution of matched substrings or the count of certain words and thus require assembled genomes to produce accurate results. These methods will not work with high accuracy when both the query and the reference are simply a set of reads. Several assembly-free methods also exist. Co-phylog [39] makes micro-alignments and calculates distances to reconstruct phylogenetic trees; Mash [45] computes the Jaccard index and an evolutionary distance using the k-mers; Simka [46] computes several distance measures based on the whole k-mer content of reads. However, these methods all assume high coverage, enough to cover most of the genome with at least one read. These levels of coverage are currently not economically feasible for building up large reference databases or for obtaining many query samples. Among existing methods, AAF [33] is the only one that aims to work even at lower coverage. AAF first infers a phylogeny and then corrects its branch lengths to reflect a given estimate of the coverage.

Here, we show that high levels of coverage are not necessary. We focus on a distance measure defined as the proportion of mismatches between the global alignment of two genomes. The mismatch rate, called genomic distance hereafter, is useful for species identification because it reflects the evolutionary divergence between two species. We introduce a new method, Skmer, for accurately computing the genomic distance even from low coverage genome-skims. In extensive test, we show that Skmer dramatically improves estimates of genomic distance based on genome-skims and accurately places genome-skim queries on to a reference collection. This assembly-free approach can therefore be considered a viable complement to currently available DNA barcoding and genome-skimming tools.

## Results

### Skmer

We decomposed reads into fixed length oligomers (denoted *k-mers* with length *k*), a technique used by many existing alignment-free methods [41, 47]. Recall that the *Jaccard index J* is a similarity measure between any two sets (e.g. k-mer collections) defined as the size of their intersection divided by the size of their union. Ondov *et al*. describe a tool, Mash [45], in which (a) *J* is estimated efficiently using a hashing procedure; and, (b) *J* is used to estimate the genomic distance between two genomes. Mash, however, assumes sufficiently high coverage. Unfortunately, *J*, in addition to the true distance, is impacted by coverage, sequencing error, and genome length. Skmer accounts for the impact of these factors on *J*.

Skmer has two stages (Fig. 1): first we use *k*-mer frequency profiles (computed using JellyFish [48]) to estimate the amount of sequencing error and the coverage (neither of which is known) using a novel method.

**Figure 1:**
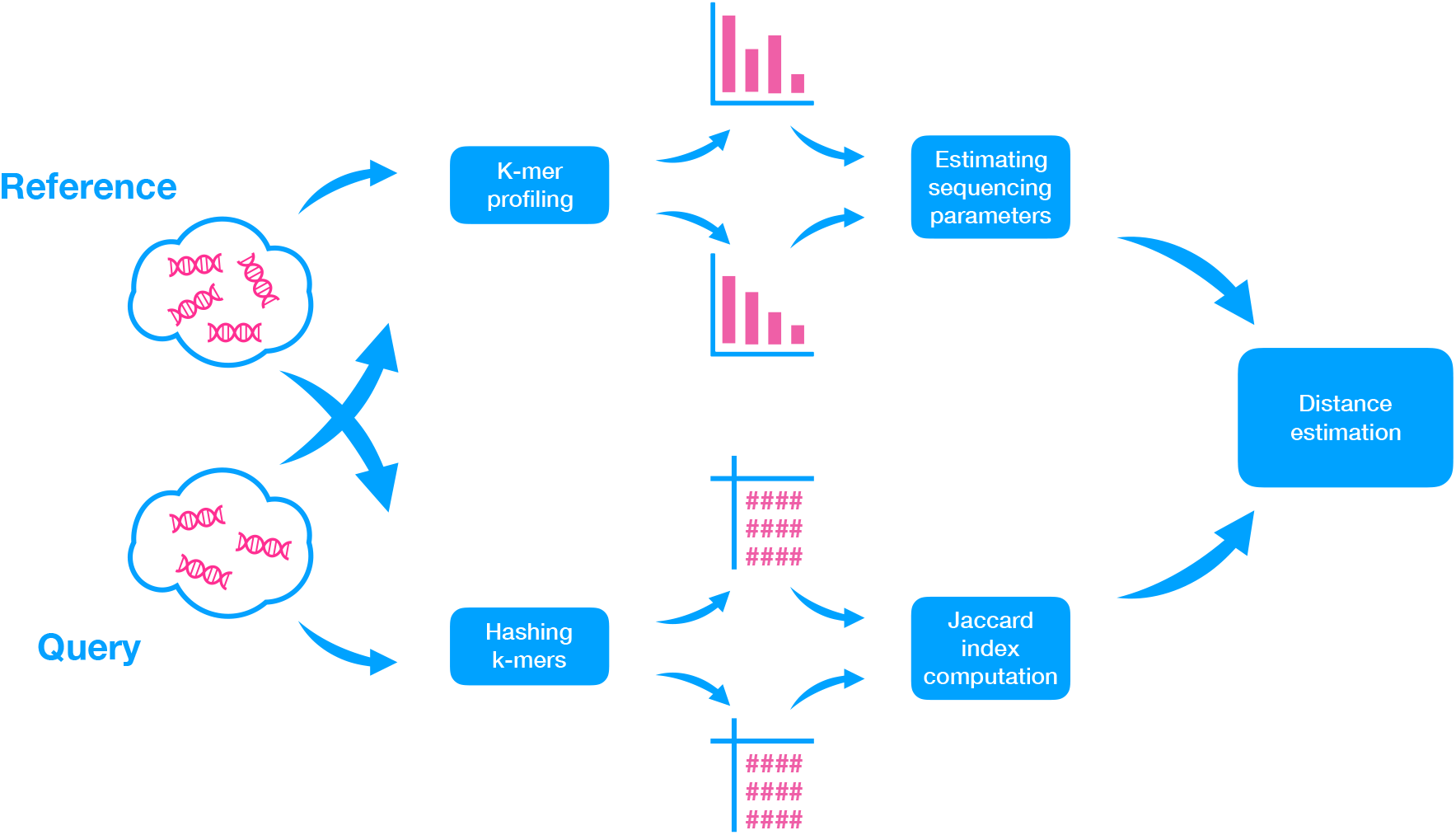
Overview of Skmer pipeline. For both query and reference genome-skims, first, the k-mer frequency profiles are used to estimate the sequencing error and coverage (top). Then, the k-mers are hashed, and a subset is retained and used to estimate the Jaccard index between the two genomes (bottom). Finally, the estimated Jaccard index and estimated sequencing coverage and error are used to compute the corrected genomic distance between the query and the reference.

Let *M_i_* be the number of *k*-mers observed *i* times in the genome-skim. Let *h* = argmax_*i*≥2_ *M_i_*. Then, defining 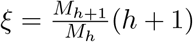, we derive (see Methods):

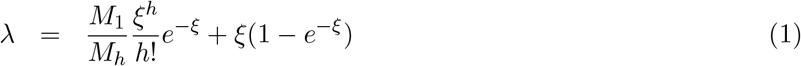

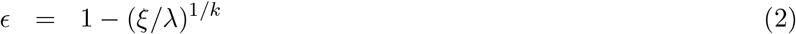

where λ and *ϵ* are our estimates of the *k*-mer coverage and the sequencing error rate, respectively.

In stage two, we use the hashing technique of Mash to compute *J*. Finally, given these estimates, we compute the genomic distance using

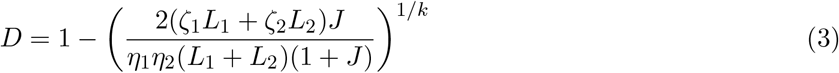

where for *i* ∈ {1, 2}, *η_i_* = 1 — *e*^−λ_*i*_(1-*ϵ_i_*)^*k*^^ and *ζ_i_* = *η_i_* + λ_*i*_(1 – (1 – *ϵ_i_*)^*k*^) (for high coverage, we define *ζ_i_* and *η_i_* differently; see Methods for details), and *L_i_* is the estimated genome length.

We used a series of experiments to study the accuracy of Skmer compared to existing methods with respect to (i) the error in computed distances, and (ii) the ability to find the closest match to a query sequence in a reference dataset of genome-skims, and (iii) phylogenetic inference. We compared the performance against *Mash* and *AAF* [33]. AAF is a method that uses *k*-mers to estimate phylogenetic distances among a set of at least four sequences. We conclude by comparing Skmer against the results of using COI barcodes from available barcode databases.

### Distance accuracy for pairs of genome-skims

We first compare the accuracy of Mash and Skmer in estimating distances between two genome skims. Since AAF outputs a phylogenetic tree and so requires at least four species, we cannot include it in our first set of analyses on pairs of genomes.

### Simulated genomes with controlled distance

Starting from the highly repetitive genome assembly of the wasp species *Cotesia vestalis*, we simulated new genomes with controlled true distance *d* by randomly adding SNPs, and then we simulated genome-skims by randomly sub-sampling reads and adding error (see Methods). On these simulated genomes, distances are computed with high accuracy by Mash when coverage is high (Fig. 2), except where the true distance is also high (i.e., 0.2). However, the accuracy of Mash quickly degrades when the coverage is reduced to 4X or less. In contrast, even when the coverage is reduced to 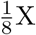, Skmer has high accuracy. For example, with the true distance set to 0.05, Mash estimates the distance as 0.081 with 1X coverage (an overestimation by 62%) while Skmer corrects the distance to 0.045 (an underestimation by 10%). Note that applying Mash* (Mash without the unnecessary approximation (1 – *D*)^*k*^ ≈ *e*^−*kD*^ used by default in Mash) to the complete assemblies generally generates very accurate results, as expected, but even given the full assembly, Mash* still has a small but noticeable error when *d* = 0.2. Note that results are extremely consistent across our ten different runs of subsampling (Fig. 2). We repeated the simulation with a lower range of coverage (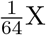 to 1X). Interestingly, even with very low coverage, the absolute distance error is small in many cases (Additional file 1: Fig. S2); however, for *d* ≥ 0.1, Skmer estimates start to degrade below 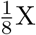 coverage.

**Figure 2:**
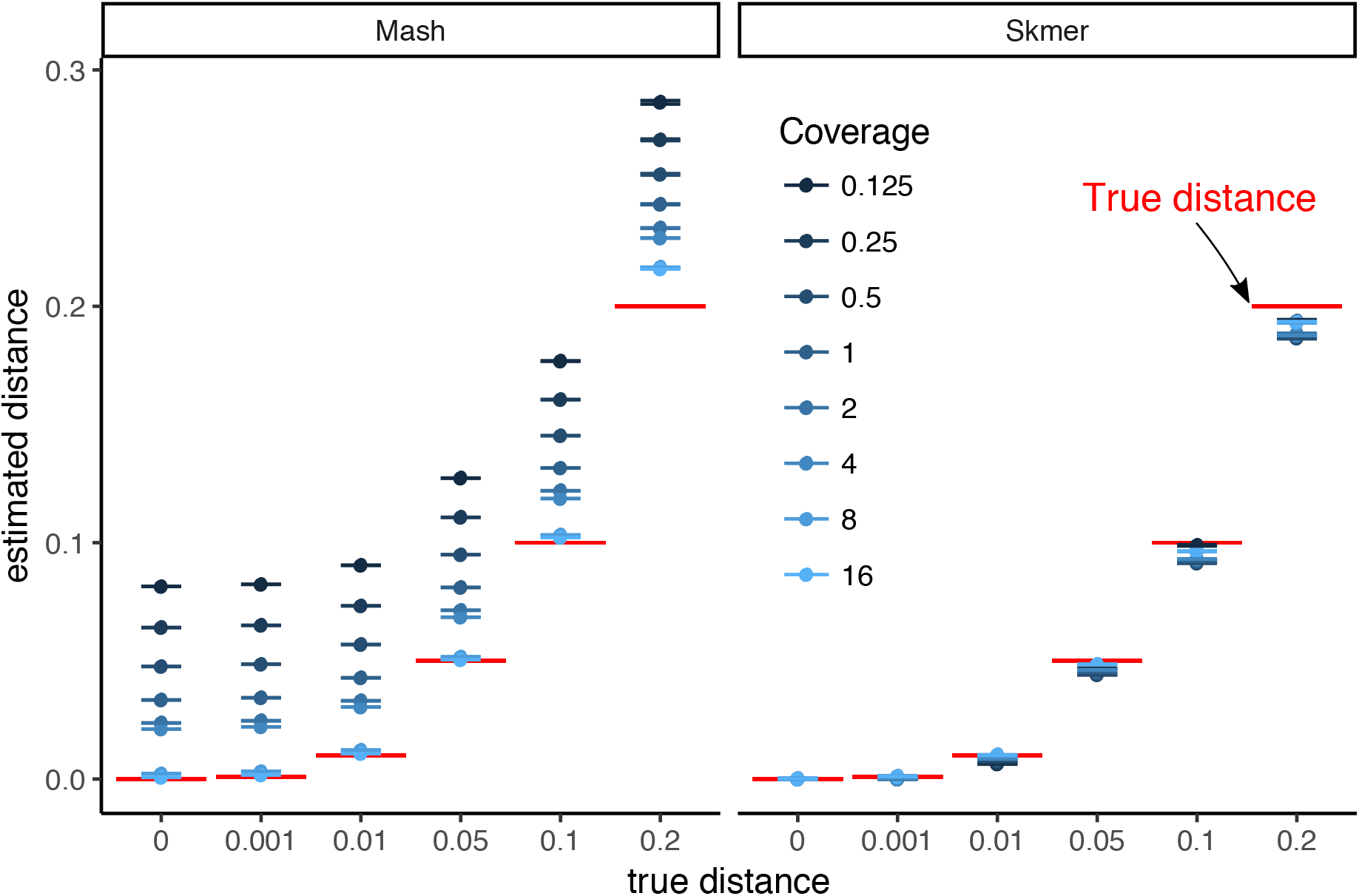
Comparing the accuracy of Mash and Skmer on simulated genomes. Genome-skims are simulated using ART with read length *ℓ* = 100. Substitutions applied to the assembly of *C. vestalis* at six different rates (x-axis), and genome-skims simulated at varying coverage range from 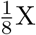 to 16X. The estimated distance (y-axis) by Mash (left) and Skmer (right) is plotted versus the real distances for each coverage level (color). The mean (dots) and standard error (lines) of distances are shown (10 repeats). True distance is shown in red. See Additional file 1: Fig. S1 for a scaled representation.

Repeating the process with the *Drosophila melanogaster* genome as the base genome also produces similar results (Additional file 1: Fig. S3). The only condition where Skmer has an absolute error larger than 0.01 is with coverage below 1X and *d* = 0.2 (Fig. 2). However, we note that for *d* = 0.001, the relative error is not small with low coverage (Additional file 1: Fig. S4b) indicating that distinguishing very small distances (perhaps below species-level) requires high coverage. Estimating the right order of magnitude when the true distance is 0.001 seems to require 2X coverage (preferably 8x) while 1X coverage is sufficient to distinguish distances at or above 0.01 (Additional file 1: Fig. S4).

### Pairs of insect and bird genomes

We now test methods on several pairs of insect and avian genomes, subsampled to create genome-skims. Note that unlike the simulated datasets, here, genomes can undergo all types of genetic variations and complex rearrangements, and thus, do not have the same length. We carefully selected several pairs of genomes to cover a wide range of mutation distance and genome length. Here, the true genomic distance is not known, but we use the distance estimated by Mash* on the full assemblies as the true distance *d*. For all pairs of insect and avian genomes (Fig. 3), Mash has high error for coverage below 8X while Skmer successfully corrects the estimated distance and obtains values extremely close to the results of running Mash* on the full assembly. For example, the distance between *A. stephensi* with length ~196Mbp and *A. maculatus* with length ~132Mbp is estimated to be 0.104 based on the full assembly and 0.102 (2% underestimation) with only 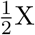 coverage using Skmer, while Mash would estimate the distance to be 0.163 (~57% overestimation).

**Figure 3:**
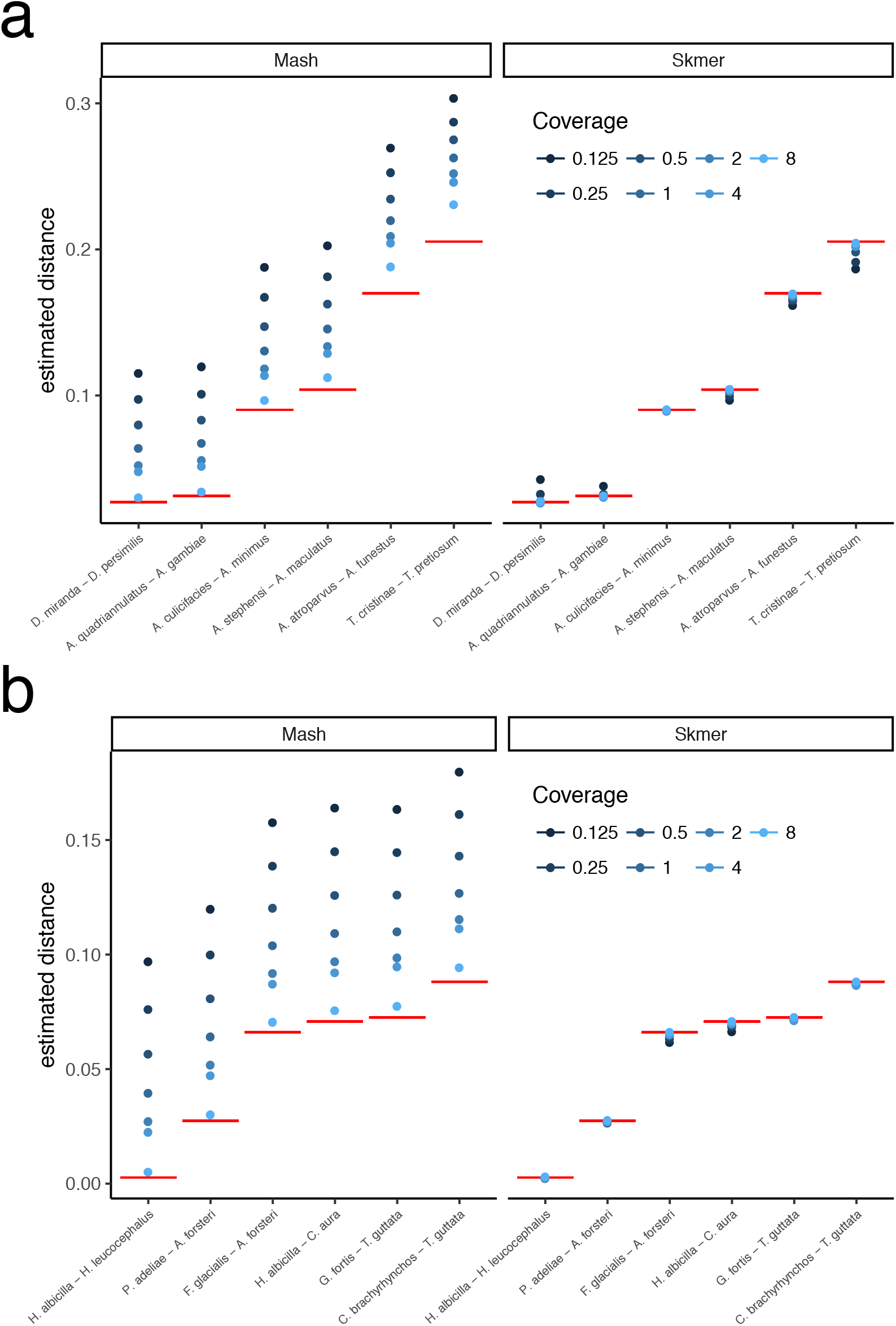
Comparing the accuracy of Mash and Skmer on pairs of insects (a) and birds (b) genomes. Genome-skims are simulated at coverage 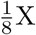 to 8X (shades of blue). The estimated distance (y-axis) is plotted for Mash (left) and Skmer (right) for each pair of species (x-axis). The results of Mash* run on assemblies, which is taken as the ground truth, is shown in red. Mash overestimates at lower coverages. Skmer estimates are closer to the ground truth and are less sensitive to the coverage. See also Additional file 1: Fig. S5.

### Distance accuracy for all pairs genome-skims

We now turn to datasets with sets of genome-skims, evaluating the accuracy of all pairs of distances. Here, since we have at least four sequences in each test, in addition to Mash, we also compare our results with AAF.

### Fixed sequencing effort

So far, our experiments have controlled for the coverage by subsampling varying amount of sequence data, proportional to the genome length. In our genome-skimming application, coverage will not be fixed. Often, the amount of sequence data obtained for each species will be relatively similar. As a result, genomes of different length end up being sequenced with different coverage depth proportional to the inverse of their length. We therefore performed a study where all species are subsampled to produce 100Mb of sequence data in total resulting in varying levels of coverage (based on the genome length, Additional file 1: Table S7). The error in the distance estimated by Mash relative to the ground truth can be quite large (higher than 300% in the worst case) while Skmer consistently makes accurate estimates close to the true distance even at the lowest amount of coverage (Fig. 4, Figs. 5, and Additional file 1: Table S8). Repeating the analysis with 0.5Gb or 1Gb total sequence data produced similar patterns, but as expected, increasing the sequencing effort reduces the error for all methods (Additional file 1: Figs. S6–S8).

**Figure 4:**
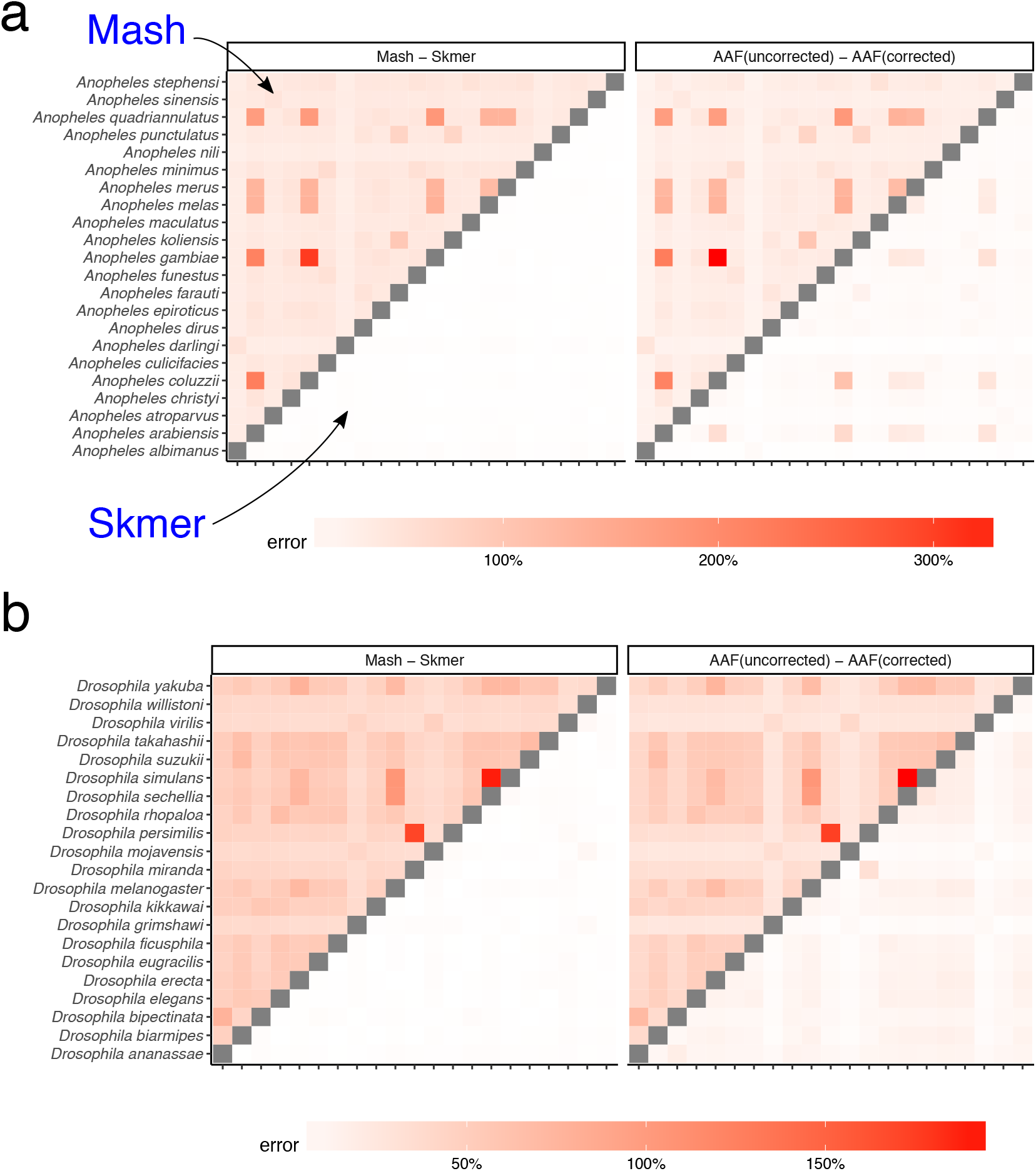
Distance error with fixed 100Mb sequence per genome for (a) 22 Anopheles, (b) 21 Drosophila. Each genome is skimmed with 100Mb sequence and distances are computed using Mash, Skmer, and AAF. True distance used in calculating the error is computed by applying each method (AAF and Mash) to the full genome assemblies. The heatmaps on the left show the error of Mash (upper triangle) and Skmer (lower triangle), and the heatmaps on the right are for AAF before correction (upper) and after correction (lower).

Before error correction, AAF has error levels that are comparable to Mash (Figs. 4b, Fig. 5b). The correction applied by AAF, similar to Skmer, reduces the negative impact of low coverage but not to the same extent. Thus, Skmer has less error compared to corrected AAF (with 100Mb sequence and across all datasets, the mean error of Skmer is 3.13% and AAF-corrected is 22.7%). For example, in the *Drosophila*, dataset, the worst-case error of AAF between any two pairs of genome-skims is 31%, whereas the error never exceeds 8% for Skmer. Note that when computing the error of AAF, we use the result of running AAF on full assemblies as the ground truth.

**Figure 5:**
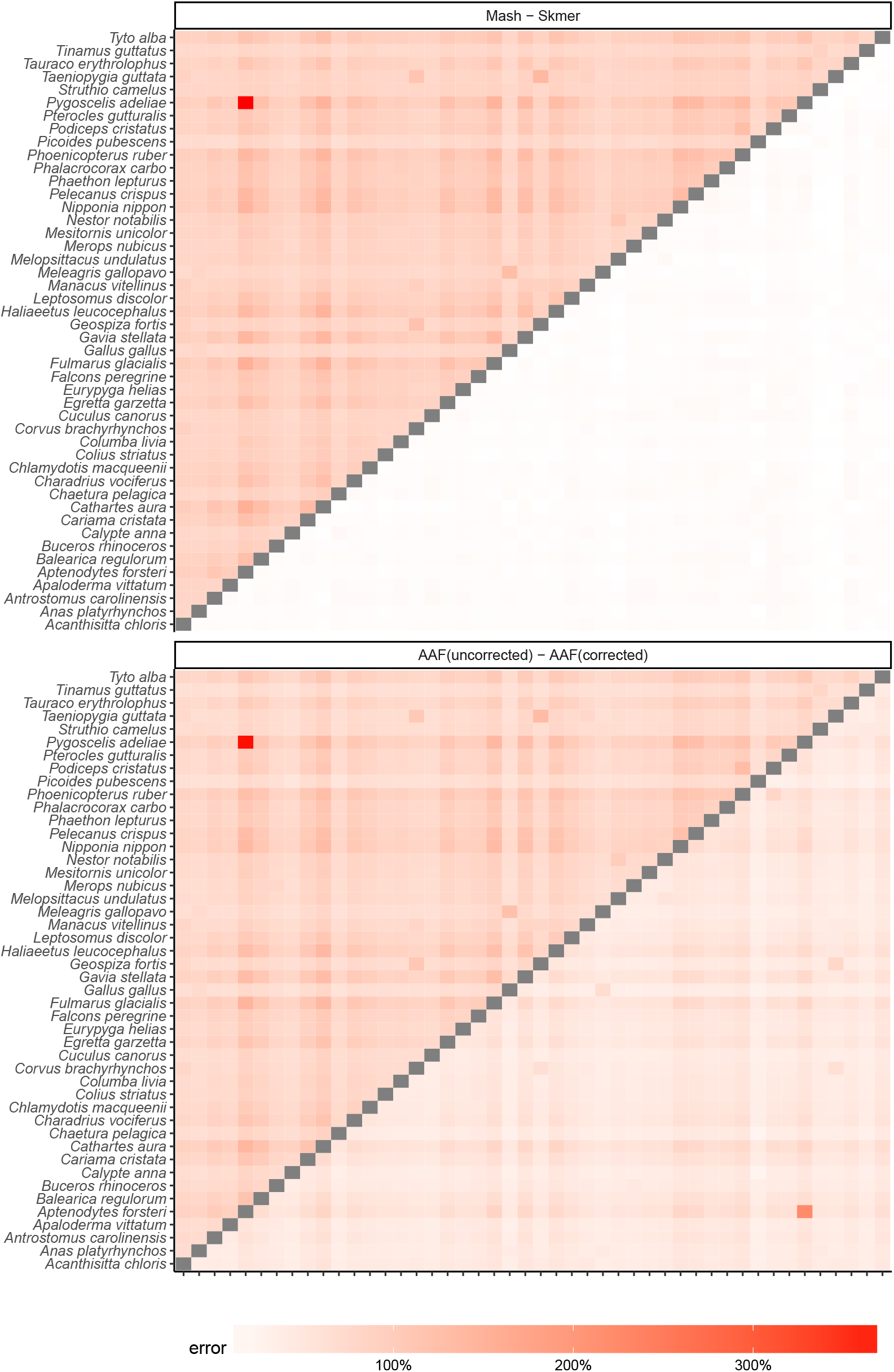
Distance error with fixed 100Mb sequence per genome for the avian dataset. The errors of Mash and AAF for the two eagle species (*H. albicilla* and *H. leucocephalus*) were extremely large (Mash: ≈ 4000%, AAF > 3000% error), dominating the color spectrum; we excluded *H. albicilla* to help readability; for the eagles, Skmer’s estimate is 0.00244 (~ 9% error).

To quantify the impact of distance estimates on downstream analyses, we used FastME [49] to infer phylogenetic trees using distances computed by Mash and Skmer on genome skims and with correction using the JC69 model [50]. AAF by default generates trees as part of its output. We compare these trees to those computed by Mash/AAF run on the full assemblies (taken as the ground truth) using the weighted Roubinson-Foulds (WRF) distance [51] (Table 1). WRF is the sum of branch length differences between the two trees (using zero length for missing branches), and we normalized WRF by the sum of branch lengths of both trees. In all three datasets, Skmer distances lead to trees with lower WRF distance to the ground truth compared to Mash and AAF/uncorrected. AAF correction reduces WRF compared to uncorrected AAF; however, Skmer trees have two to 14 times less error compared to the corrected AAF, except in one case where AAF/corrected has 1.05% error and Skmer has 1.19% (Table 1). Increasing the size of skims to 0.5Gb and 1Gb helps all methods to produce more accurate trees.

**Table 1:** Tree error. For each method, we show normalized weighted RF distance (%) of trees inferred from genome-skim distances to trees inferred from full assembly distances. Boldface: the lowest error.

| Dataset | Sequencing effort | Mash | Skmer | AAF (uncorrected) | AAF (corrected) |
| --- | --- | --- | --- | --- | --- |
| Anopheles | 0.1G | 23.19% | **1.07%** | 19.92% | 6.36% |
|  | 0.5G | 12.84% | **0.45%** | 9.74% | 4.9% |
|  | 1G | 8.92% | **0.37%** | 9.59% | 3.3% |
|  | Mixed | 14.75% | **0.58%** | 8.46% | 8.45% |
| Drosophila | 0.1G | 23.87% | **2.05%** | 20.29% | 5.85% |
|  | 0.5G | 13.33% | **0.72%** | 10.37% | 5.25% |
|  | 1G | 7.11% | **0.58%** | 10.84% | 2.2% |
|  | Mixed | 16.58% | **1.11%** | 11.36% | 10.87% |
| Birds | 0.1G | 37.03% | **5.64%** | 31.81% | 21.13% |
|  | 0.5G | 25.16% | **1.91%** | 20.8% | 6.86% |
|  | 1G | 19.42% | 1.19% | 15.54% | **1.05%** |
|  | Mixed | 28.14% | **3.08%** | 18.15% | 7.57% |

### Heterogeneous sequencing effort

In addition to changes in the genomic length, the sequencing effort per species may also vary across sequencing protocols, experiments and research labs, and so a database of reference genome-skims may consist of samples with heterogeneous sequencing efforts. To capture this, for each species, we choose its total sequencing effort from three possible values 0.1Gb, 0.5Gb, and 1Gb, uniformly at random, and estimate all pairs of distances within each dataset as before (Fig. 6 and Additional file 1: Fig. S9). Similar to the case of fixed sequencing effort, Skmer mitigates large relative error in the distances estimated by Mash and produces more accurate results than both Mash and AAF, (Table 2, Fig. 6, and Additional file 1: Fig. S9). For example, comparing to the case of fixed 100Mb genome-skims of the *Drosophila* dataset, the worst-case error of AAF is increased to 70%, while using Skmer it remains almost the same (8%). Comparing trees inferred from distances estimated by various methods also confirms the higher accuracy of Skmer (Table 1). For instance, on the Anopheles dataset, Skmer has only 0.58% WRF distance to the reference tree whereas Mash and AAF-corrected trees have 14.75% and 8.45% WRF distance.

**Figure 6:**
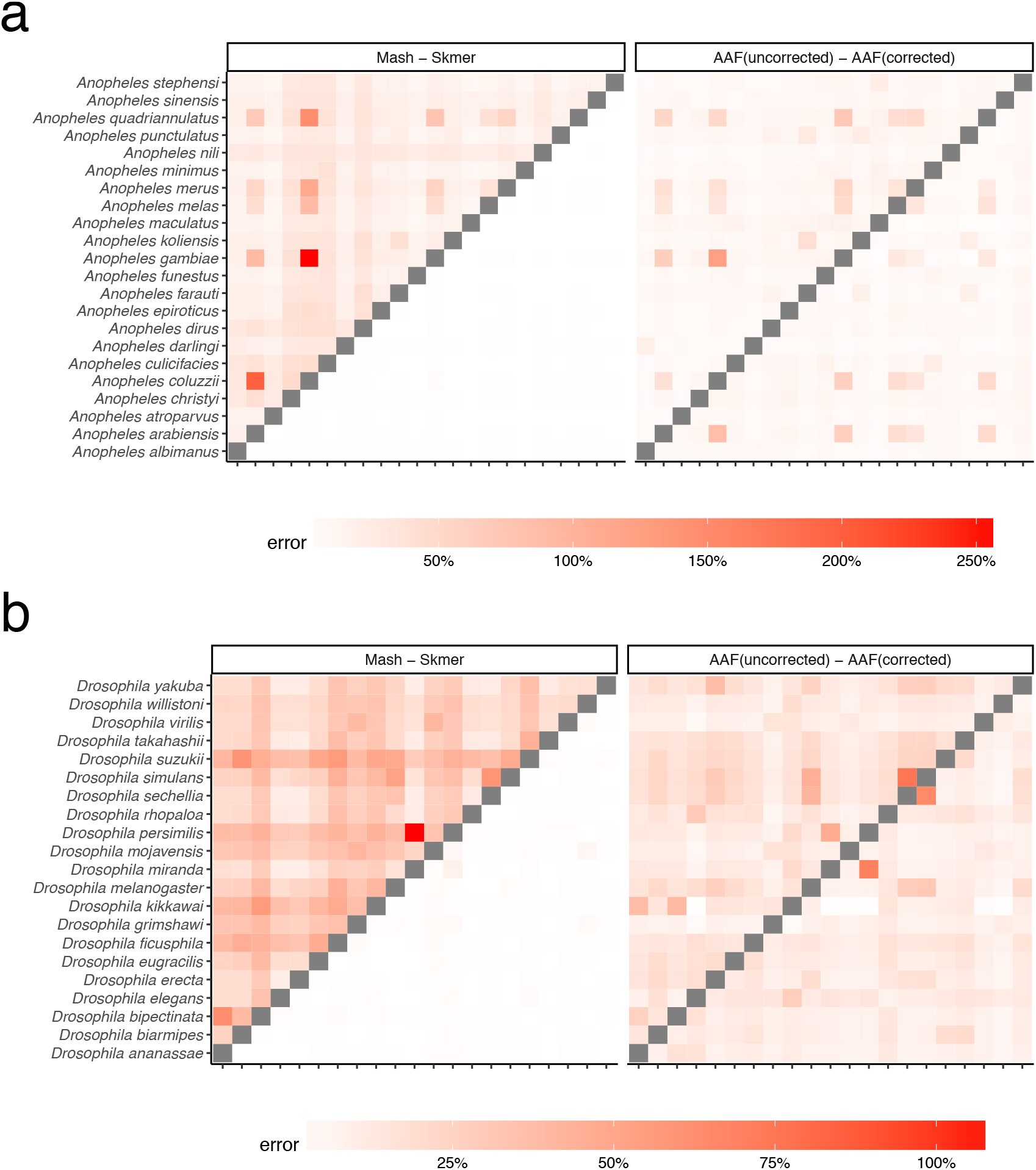
Distance error with heterogeneous sequencing effort for (a) Anopheles and (b) Drosophila. Species have random amount of sequence chosen uniformly among 0.1Gb, 0.5Gb, and 1Gb. See Additional file 1: Fig. S9 for birds.

**Figure 7:**
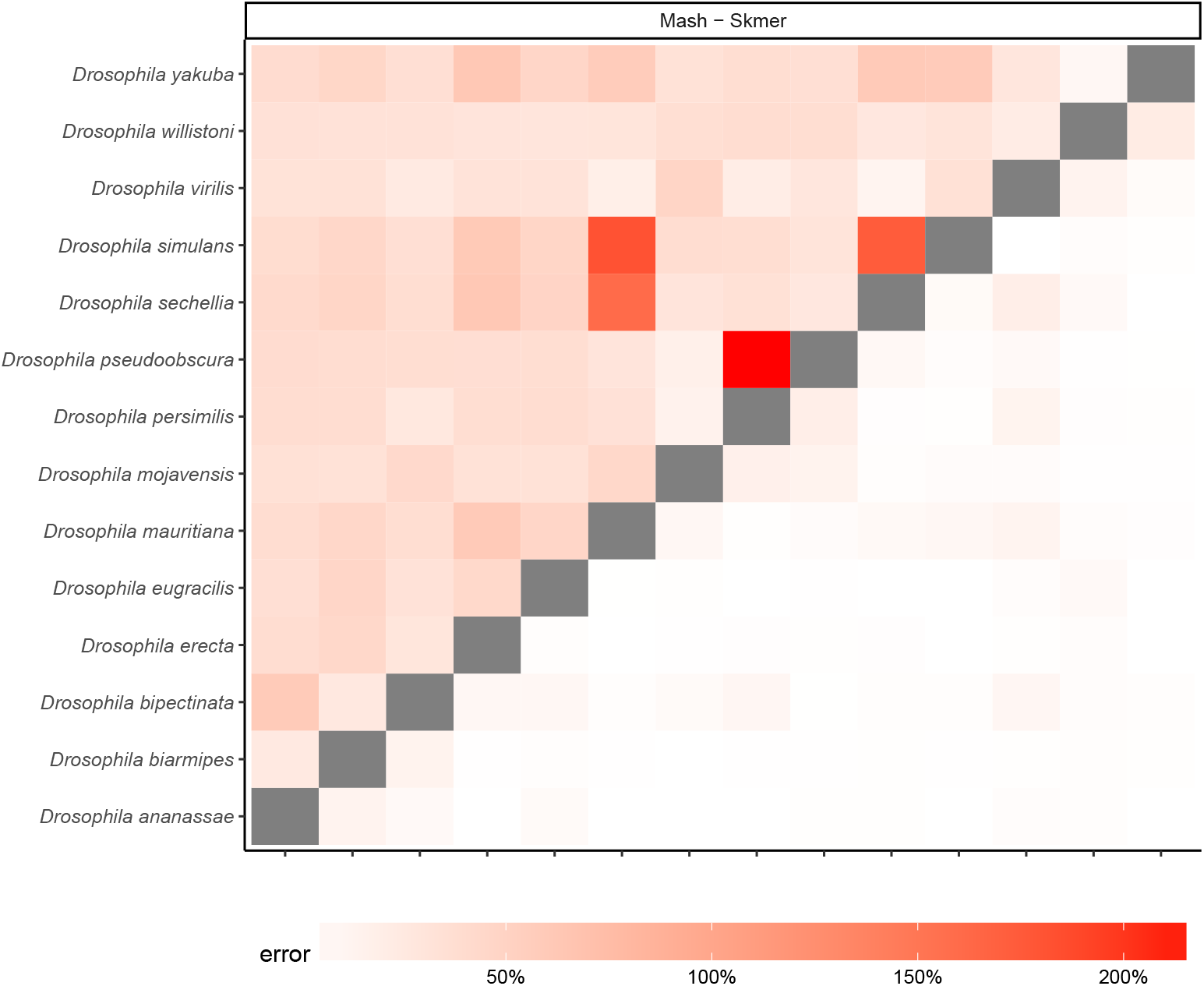
Comparing the error of Mash and Skmer on a dataset of 14 Drosophila genome-skims. Each SRA is subsampled to 100Mb and then filtered to remove contamination. True distances are computed from the assemblies.

**Figure 8:**
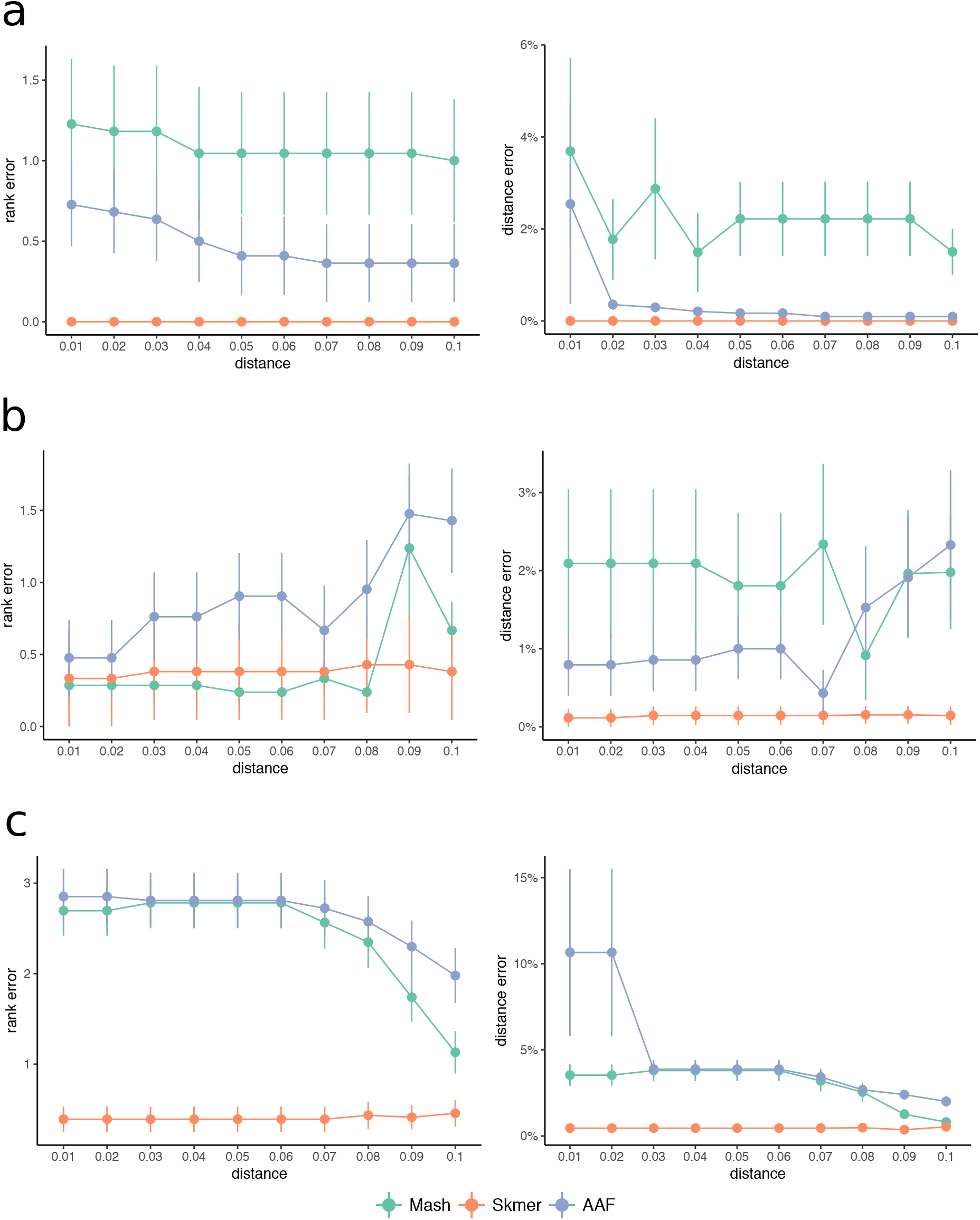
The mean rank and distance error of the best remaining match in leave-out experiments. The distance of closest genome in the reference to a query is varied from 0.01 to 0.1 (x-axis). The rank and distance errors (y-axis) of the best match to a query, are computed by comparing the order given by each method with the order obtained by applying Mash* to the full assemblies (ground truth). For each dataset, the experiment is repeated by taking each species as the query, and then the errors are averaged. Three methods, Mash, Skmer, and AAF, are compared on: (**a**) the *Anopheles* dataset, (**b**) the *Drosophila* dataset, and (**c**) the avian dataset.

**Table 2:** Comparing the average error of Mash, Skmer, and AAF in estimating distances over three datasets with heterogeneous sequencing effort.

| Dataset | Mash | Skmer | AAF (uncorrected) | AAF (corrected) |
| --- | --- | --- | --- | --- |
| *Anopheles* | 28.72% (1.10%) | **0.84%** (0.03%) | 13.48% (0.56%) | 11.36% (0.44%) |
| *Drosophila* | 29.05% (0.59%) | **0.84%** (0.04%) | 15.25% (0.38%) | 10.94% (0.33%) |
| Birds | 64.29% (0.54%) | **2.21%** (0.04%) | 36.02% (0.29%) | 5.28% (0.16%) |

\* The standard error of the mean is provided in parentheses.

### Genome skims from real reads

So far, all of our tests used simulated reads. When analyzing real genome skims, there are additional complications such as extraneous DNA (real or artifactual) and the over representation of organelle genome. We next tested Skmer using real reads. We created 100Mb skims of 14 Drosophila genomes by subsampling short-read data produced in a recent Drosophila genome assembly study [52]. Before running Skmer or Mash, we filtered reads that (even partially) aligned to 12 Drosophila-associated microbial genomes as reported in previous studies [53–55] (see Table S3), to the human genome, or to the mitochondrial genome of respective Drosophila species. We then estimated all pairs of distances as before and computed the error relative to the distances computed from the assemblies (Fig 7). Consistent with the results we obtained on the simulated skims, Skmer has less error compared to Mash. The average error of Mash on this dataset is 43.48% (± 2.29%) with maximum error of 217%. Skmer, on the other hand, has an average error of 4.21% (± 0.35%) and its maximum error is 22.2%.

### Running time

Skmer and Mash have comparable running time, while AAF is much slower. In the experiment with heterogeneous sequencing effort, the total running time (using 24 CPU cores) to compute distances based on genome-skims for all 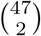 pairs of birds using Mash, Skmer, and AAF was roughly 8, 33, and 460 minutes, respectively.

### Leave-out search against a reference database of genome-skims

We now study the effectiveness of using genomic distance to search a database of genome-skims to find the closest match to a query genome-skim. Given a query genome-skim and a reference dataset of genomes, we can order the reference genomes based on their distance to the query. The results can be provided to the user as a ranking. When the query genome is available in the reference dataset, finding the match is relatively easy. To study the effectiveness of the search as the distance of the closest available match increases, we use a leave-out experiment, as described in Methods. Figure 8 shows the mean rank error as well as the mean distance error of the best remaining match in a leave-out experiment when removing genomes closer than *d* for 0.01 ≤ *d* ≤ 0.1. A rank error (or distance error) equal to zero corresponds to a perfect match to the best available genome.

On all three datasets, Skmer consistently and often substantially outperforms Mash and AAF in terms of finding the best remaining match, except the *Drosophila* dataset where Mash and Skmer have comparable rank error, while both are better than AAF (Fig 8). Even in that case, on average, the distance of the best match found by Skmer is closer to the distance of the true best match compared to the best hit found by Mash. Moreover, the mean rank error of Skmer is smaller than Mash (Additional file 1: Fig. S10) if we exclude only one species *Drosophila willistoni* (which is at distance 0.1565 ≤ *d* ≤ 0.1622 from other species). It is also notable that over the avian dataset, Skmer has mean rank error less than 0.5 for all range of distances, while Mash and AAF can be off by more than 2.5 on average. These results demonstrate that correcting the distance not only impacts our understanding of the absolute distance, but also, impacts results of searching a reference library.

### Phylogeny reconstruction and comparison to organelle markers

As the last experiment, we estimated phylogenetic trees for *Anopheles* and *Drosophila* datasets after transforming the genomic distances estimated by Skmer to Jukes-Cantor (JC) distances [50]. For each dataset, we also built another tree based on available COI barcodes, using an identical method. We compare the results against a reference tree obtained from Open Tree of Life [56]. We restricted the results to species for which COI barcodes were available (Fig. 9ab).

**Figure 9:**
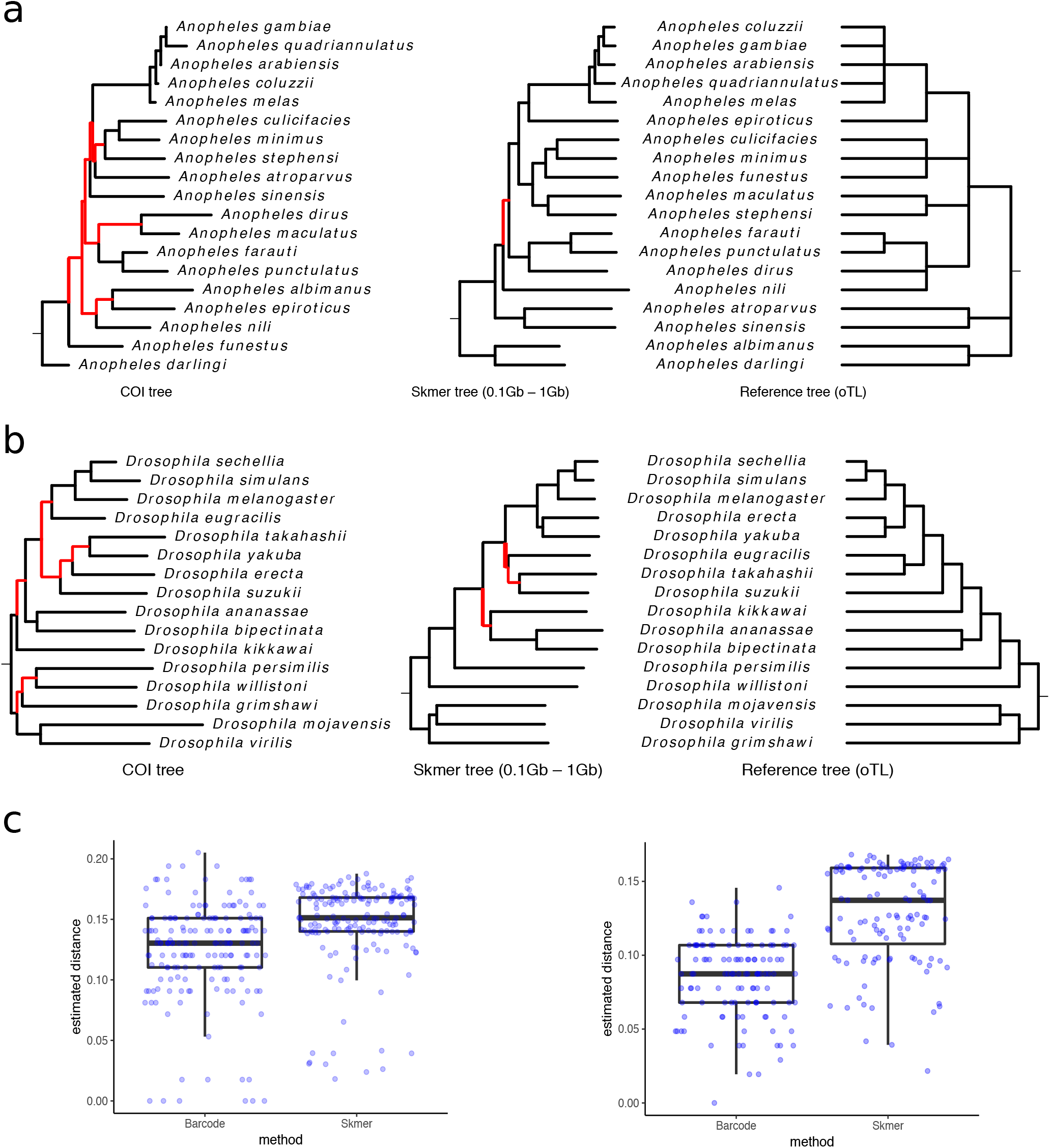
Comparing distances and phylogenetic trees from COI barcodes and simulated genome-skims. Shown in red are wrong internal branches corresponding to the bipartitions that are not found in the reference tree. Genome-skim size is randomly chosen among 0.1Gb, 0.5Gb, and 1Gb. (**a**) *Anopheles* trees. (**b**) *Drosophila* trees. (**c**) Distribution of distances for *Anopheles* (left) and *Drosophila*, (right) genomes

For the *Anopheles* species, Skmer distances produce a tree that is almost identical to the reference tree (with only one branch difference out of nine), while COI tree differs from the reference in seven branches. Similarly, for the *Drosophila* species, Skmer differs from the reference in three branches (with small local changes) out of 13 total branches in the reference tree, whereas COI tree is very inconsistent with the reference tree (seven branches are different). We also built maximum-likelihood trees from COI barcodes (Additional file 1: Fig. S11), but the number of incorrect branches did not reduce. Comparing the distribution of all pairwise genomic distances obtained from genome-skims and barcodes (Fig. 9c), Skmer has larger distances and fewer pairs with zero or close to zero distance, indicating that Skmer has a higher resolution in differentiating between samples. For example, four species of the *Anopheles* genus *A. coluzzii*, *A. gambiae*, *A. arabiensis*, and *A. melas* have very small pairwise distances based on COI barcodes, while using Skmer, the estimated distances are in the range 0.02–0.04 for these species.

## Discussion

We showed that Skmer can compute the genomic distance between a pair of species from genome-skims with very low coverage (at or even below 1X), with much better accuracy than the main two alternatives, Mash and AAF. We also showed that the distances computed by Skmer can accurately place a voucher genome-skim within a reference database of genome-skims, and can be used to infer the phylogenetic tree with reasonable accuracy. While Skmer is not the first *k*-mer based approach for distance estimation or phylogenetic reconstruction, as we showed, the alternatives have low accuracy given low coverage data. We compare with Mash because it is used within Skmer and is one of the most widely-used alignment and assembly-free methods. However, we note that authors of Mash do no claim it can handle low coverage, and so our results are not a criticism of their approach. Besides the methods we discussed, many other alignment-free sequence comparison and phylogeny reconstruction algorithms exist [25, 28, 29, 31, 32, 34–43]. However, these methods take as input assembled (but unaligned) sequences, and thus, are not applicable in an assembly-free pipeline. In other words, their goal, is to avoid the alignment step and not the assembly step.

Compared to using COI markers, currently used in practice, we showed that using *all k*-mers, including those from the nuclear genome, improves the phylogenetic accuracy. These improvements are resulting from distances that have a larger range and more resolution compared to COI. Also, the increased resolution should not be surprising given that the entire genome is much larger than any single locus, reducing the variance in estimates of the distance. Beyond the question of resolution, gene trees and species trees need not match [57], a fact that can further reduce the accuracy of marker genes for both species identification and phylogeny reconstruction. By using the entire genome, Skmer ensures that an average distance across the genome is computed, reducing the sensitivity to gene tree/species tree discordances. Moreover, a recent result shows that the JC-transformed genomic distance is a statistically consistent estimator of the species distances despite gene tree discordance due to incomplete lineage sorting [58], further encouraging our use of the genomic distance as a measure of the evolutionary divergence.

We showed that genomic distances as small as 0.01 can be estimated accurately from genome-skims with 1X or lower coverage. What does a distance of 0.01 mean? The answer will depend on the organisms of interest. For example, two eagle species of the same genus (*H. albicilla* and *H. leucocephalus*) have *D* ≈ 0.003 but two *Anopheles* species of the same species complex (*A. gambiae* and *A. coluzzii*) have *D* ≈ 0.018. Broadly speaking, for eukaryotes, detecting distances in the 10^-2^ order is often enough to distinguish between species (Additional file 1: Fig. S12). On the other hand, to differentiate individuals in a population, or very similar species, we may need to reliably estimate distances of the order 10^-3^. Detection at these lower levels seems to require > 1X coverage using Skmer (Additional file 1: Fig. S4b) but future work should study the exact level of sequencing required for accurate ordering of species at distances in the order of 10^-3^ or less. Moreover, the question of the minimum coverage required may avail itself to information-theoretical bounds and near-optimal solutions, similar to those established for the assembly problem [59, 60].

Although most of our tests simulated genome skims simulated from assemblies, we also tested Skmer on genome skims simulated by subsampling previous whole genome sequencing experiments. Several complications have to be addressed in real applications. The actual coverage of real genome skims may not be uniform and randomly distributed and they can have an overrepresentation of mitochondrial or plastid sequence. More importantly, other sources of DNA originating from for example, parasites, diet, fungi, commensals, bacteria, and human contamination may all be present in the sample and may cause a bias in the estimation of distances. In our test, we simply searched all reads in a genome-skim against a few bacterial genomes and the human reference genome; this simple scheme filtered out up to ~10% of reads (for *D. virilis*). These filtering strategies were sufficient to produce reliable distance estimates in the case of Drosophila genomes. We recommend that before using Skmer, such database searches should be used to find and eliminate bacterial or fungal contamination (using BLAST [61] or perhaps metagenomic tools such as Kraken [62]), as well as removing contaminant reads with human origin (using for example Bowtie2 [63]). However, in future, it will be beneficial to develop better methods for finding extraneous reads without reliance on known sources.

A related direction of future work is to explore whether Skmer can be extended to environmental DNA analyses, i.e., queries consisting of genome-skims of multi-taxa samples. While Skmer is presented here in a general setting, its best use is for eukaryotic organisms, where the notion of species is better established and species can be separated with reasonable effort. We tested Skmer on birds and insects, but we predict it will work equally well for plants, a prediction that we plan to test in future work.

Throughout our experiments, we used Mash* run on the assemblies to compute the ground truth. Given the true alignment of the two genomes, we can compute the true genomic distance as the proportion of mismatches among *aligned* orthologous positions (i.e., ignoring gaps). To ensure that Mash* closely approximates true distances, we used simulated genomes of Rat and Mouse from the Mammalian dataset of the Alignathon competition [64]. This simulation uses Evolver [65] and includes many forms of mutation, including indels, rearrangement, duplications, and losses. On this dataset, the true distance based on the known true alignment is 0.145 and Mash* estimated the distance as 0.143, which is a very good approximation. In contrast, FastANI [66], an alignment-free sequence mapping tool for estimating average nucleotide identity, computes the distance as 0.189. If we count gaps as non-matching positions in the definition of distance, then the true distance would be 0.287, which also does not match FastANI. Presumably, FastANI, which relies on alignment of short blocks, counts short gaps (with *some* definition of short) as mismatch but excludes larger ones. Thus, on real data, Mash* is the best available option to approximate the true distance. Finally, note that, for real genomes, we chose not to use estimated whole genome alignments (WGA) to compute the ground truth because WGA is a difficult problem, and WGAs that are available are not necessarily accurate. We get inconsistent estimates of distance when we use pairwise or multiple WGAs. For example, between *D. melanogaster* and *D. yakuba*, the distance changes from 0.10 when using the multiple WGA [67], to 0.21 if we use the pairwise WGAs [68] from the UCSC genome browser [69], which is the state-of-the-art.

The connection between genomic distance and phylogenetic distance depends on mutation processes considered. If only substitutions are allowed and assuming the Jukes-Cantor model, the phylogenetic distance is 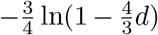; note this transformation is monotonic and does not change rankings of matches to a query search. Assuming a more complex model such as GTR [70], genomic distance is not enough to estimate the phylogenetic distance. However, we have devised a simple procedure to estimate GTR distances using the log-det approach [71] by repeated applications of Skmer to perturbed reads (Additional file 1: Appendix B). The GTR distances can rank matches to a query differently from the genomic distance; the accuracy of the two distances should be compared in future work.

Insertions, deletions, duplications, losses, and repeats can all lead to differences between genomes, thereby reducing the Jaccard index and increasing the genomic distance. They also impact genomic length. Interestingly, in our experiments, Skmer run with the true coverage is *less* accurate than with estimated coverage (Additional file 1: Fig. S13). We speculate that on genomes with repeats, by overestimating coverage, our method gives an estimate of the “ effective” coverage, reducing the impact of repeats on the Jaccard index. Nevertheless, with these complex mutations, the correct definitions of the evolutionary distance and genomic distance are not straightforward; nor is it clear how the Jaccard index should be translated to the genomic distance. Here, we used a heuristic approach that simply averaged the length of the two genome, leaving these broader questions about the best definition of genomic distance in the presence of large structural variations to future work.

## Conclusions

Skmer is an assembly-free and alignment-free tool for estimating the distance between two genome-skims. It can estimate a wide-range of distances with high accuracy from low-coverage and mixed-coverage genome-skims with no prior knowledge of the coverage or the sequencing error. Our paper shows that the idea of genome-wide sample identification using genome-skims has merit and should be pursued in the future.

## Methods

Consider an idealized model where two genomes are the outcome of a random process that copies a genome and introduces mutations at each position with fixed probability *d*. Moreover, substitutions are the only allowed mutation. In this case, the per-nucleotide hamming distance *D* between the two genomes is a random variable (r.v.) with expected value *d*. We would like to estimate *d*. While this is a simplified model, we will test the method on real pairs of genomes that differ due to complex mutational processes (also, see Additional file 1: Appendix B for extensions). We start with known results connecting the Jaccard index and the hamming distance and then show how these results can be generalized to low coverage genome-skims. Throughout, we present our results succinctly and present derivations and more careful justifications in Additional file 1: Appendix A of the supplementary material.

### Jaccard index versus genomic distance

The Jaccard index of subsets *A*_1_ and *A*_2_ is defined as

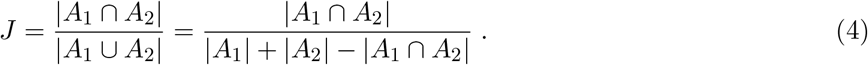

Let *W* be the number of shared *k*-mers between the two genomes. Note that: 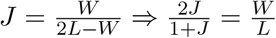, where *L* is the genome length. Assuming random genomes and no repeats, perhaps justifiably [72], the probability that a changed *k*-mer exists elsewhere in the genome is vanishingly small for sufficiently large *k*. Thus, we assume a *k*-mer is in the shared *k*-mers set only if no mutation falls on it, an event that has probability (1 – *d*)^*k*^. Thus, we can model *W* as a binomial with probability (1 – *d*)^*k*^ and *L* trials. As Ondov *et al*. [45] pointed out, we can estimate

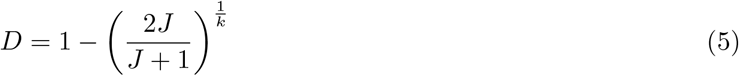

and they further approximate *D* as 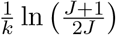. To be able to estimate large distances, we avoid the unnecessary approximation and use Equation 5 directly. We skim each genome to obtain *k*-mer sets *A*_1_, *A*_2_ and estimate *J* using Equation 4, which can be computed efficiently using a hashing technique used by Mash [45]. Note that, however, Equation 5 assumes a high coverage of the genome so that each *k*-mer is sampled at least once with very high probability. This assumption is violated for genome-skims in consequential ways. As a simple example, suppose the coverage is low enough that a *k*-mer is sampled with probability 0.5. Then, even for identical genomes, we estimate *J* as 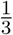, resulting in a distance estimate of *D* ≈ 0.032 for *k* = 21.

### Extending to genome-skims with known low coverage and error

We now show how Equation 5 can be refined to handle genome-skims despite low and uneven coverage, sequencing error, and varying genome-lengths. We first assume that coverage and error are known and later show how to compute these.

### Low coverage

When the genome is not fully covered, three sources of randomness are at work: mutations and sampling of *k*-mers from each of the two genomes. Each genome of length *L* is sequenced independently using randomly distributed short reads of length *ℓ* at coverages *c*_1_ and *c*_2_ to produce two genome-skims. Under the simplifying assumption that genomes are not repetitive, we choose *k* to be large enough so that each *k*-mer is unique with high probability. Therefore, the number of distinct *k*-mers in each genome is *L* – *k* ≃ *L*. The probability of covering each *k*-mer can be approximated as *η_i_* = 1 – *e*^−*λ_i_*^ where *λ_i_* = *c_i_*(1 – *k*/*ℓ*). Modeling the sampling of *k*-mers as independent Bernoulli trials, |*A_i_*| becomes binomially distributed with parameters *η_i_* and *L*. By independence, *W* = |*A*_1_ ∩ *A*_2_| also becomes binomially distributed with parameters *η*_1_*η*_2_(1 – *d*)^*k*^ and *L*. Moreover, *U* = |*A*_1_ ∪ *A*_2_| can also be modeled approximately as a Gaussian with mean (*η*_1_+*η*_2_–*η*_1_*η*_2_(1–*d*)^*k*^)*L*. Treating *η*_1_ and *η*_2_ as known and dividing 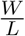 by 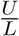 gives us:

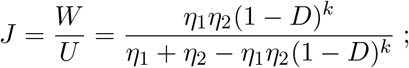

thus,

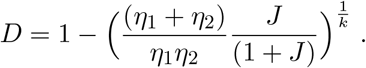

### Sequencing error

Each error reduces the number of shared *k*-mers and increases the total number of observed *k*-mers, and thus can also change the Jaccard index. Let *ϵ_i_* denote the base-miscall rate for genome skim *i*. For large *k* and small *ϵ_i_*, the probability that an erroneous *k*-mer produces a non-novel *k*-mer is negligible. The probability that a *k*-mers is covered by at least one read, without any error, is approximately

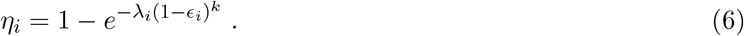

Adding up the number of error-free and erroneous *k*-mers, the total number of *k*-mers observed from both genomes can again be approximately modeled as a Gaussian with mean *ζ_i_L* for

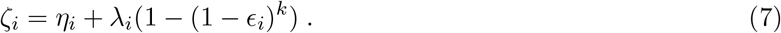

Just as before, we can simply estimate *D* by solving for it in

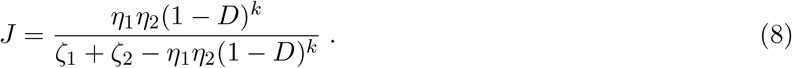

When the coverage is sufficiently high, each *k*-mer will be covered by multiple reads with high probability, and low-abundance *k*-mers can be safely considered as erroneous. Mash has an option to filter out *k*-mers with abundances less than some threshold *m* to remove *k*-mers that are likely to be erroneous. In this case,

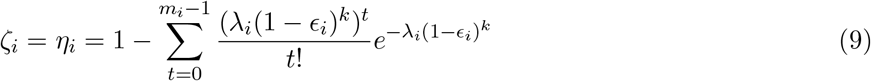

assuming all erroneous *k*-mers are removed. For instance, filtering single-copy *k*-mers (i.e., *m* = 2) gives us:

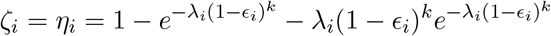

and the Jaccard index follows the same equation as (8). Since this filtering approach only works for high coverage, we filter low coverage *k*-mers only when our estimated coverage is higher than a threshold (described below). Note that the genome-skims compared may use different filtering schemes yet Eqn. 8 holds regardless.

### Differing genome lengths

Based on a model where the genomic distance between genomes of different lengths is defined to be confined to the mutations that are falling on homologous sequences, we can drive

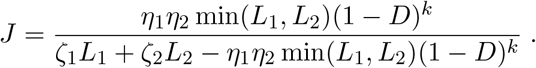

This computation does not penalize for genome length difference. While a rigorous modeling of evolutionary distance for genomes of different length require sophisticated models of gene gain, duplication, and loss, we take the heuristic approach used by Ondov *et al*. [45] and simply replace min(*L*_1_, *L*_2_) with (*L*_1_ + *L*_2_)/2. This ensures that the estimated distance increases as genome lengths becomes successively more different. This leads us to our final estimate of distance given by:

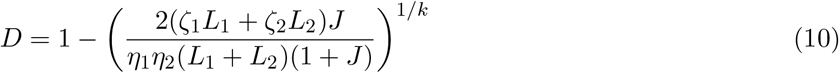

### Estimating sequencing coverage and error rate

So far we have assumed a perfect knowledge of sequencing depth and error. However, for genome-skims, the genome length is not known; thus, we need to estimate the coverage in order to apply our distance correction. We also assume a constant base error rate, and co-estimate it with the coverage.

The sequencing depth, which is the average number of reads covering a position in the genome, can be estimated from the *k*-mer coverage profiles. The probability distribution of the number of reads covering a *k*-mer is a Poisson r.v. with mean λ, where λ is defined as *k*-mer coverage. As we look into the histogram data, it is easier to work with counts instead of probabilities. Let *M* denote the total number of *k*-mers of length *k* in the genome, and *M_i_* count the number of *k*-mers covered by *i* reads. Thus, for *i* ≥ 0, 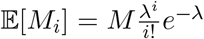. For a given set of reads, we can count the number of times that each *k*-mer is seen, and assuming zero sequencing error, it equals the number of reads covering that *k*-mer. Then, we can aggregate the number of *k*-mers covered by *i* reads and find *M_i_* for *i* ≥ 1. However, since in a genome-skim, large parts of the genome may not be covered, both *M* and *M*_0_ are unknown. To deal with this issue, we could take the ratio of consecutive counts to get a series of estimates of λ as 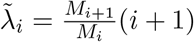 for *i* = 1, 2,.… In practice, sequencing errors change the frequency of *k*-mers and has to be considered when estimating the coverage. Assuming that the error is introduced at a constant rate along the reads, we can use the information in the *k*-mer counts to co-estimate e and *ϵ*. Like before, we assume that the *k*-mer length *k* is large enough that any error will introduce a novel *k*-mer, so the count of all erroneous *k*-mers is added to the count of single-copy *k*-mers. Moreover, for *k*-mers with more than one copy, the number of times that each kmer is seen equals the number of reads covering that *k*-mer without any error. Formally, let *M_i_* denote the count of *k*-mers seen *i* times in the presence of error, and *ρ* = (1 – *ϵ*)^*k*^ denote the probability of error-free *k*-mer.

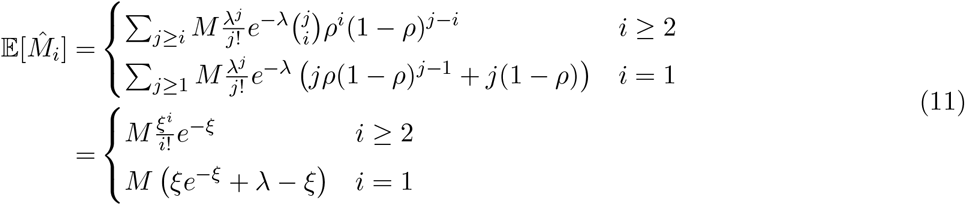

where *ξ* = *λρ* is the average number of error-free reads covering a k-mer. A family of estimates for *ξ* is obtained by taking the ratio of consecutive counts of error-free k-mers as 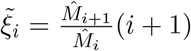 for *i* ≥ 2. Then, using an estimate of *ξ* and the count of single-copy k-mers, we get a series of estimates of λ for *i* ≥ 2 as

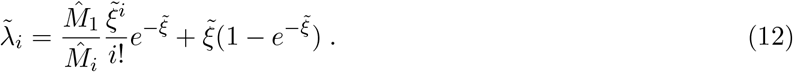

Moreover, we can estimate the error rate from the estimates of λ and *ξ* as

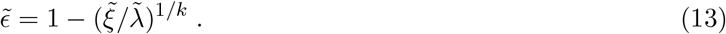

While any of these 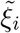 and 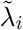 can be used in principle, the empirical performance can be affected by the choice; in our tool, we use heuristic rules (described below) that seek to use large *M_i_* values.

### Skmer: implementation

Skmer takes as input two or more genome-skims. It uses JellyFish [48] to compute *M_i_* values, which are then used in estimating λ and *ϵ* based on Equations 12 and 13, by setting 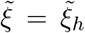 and 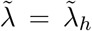, where *h* = argmax_*i*≥2_ *M_i_*. Then, Mash is used to estimate the Jaccard index, with *k* = 31 (selected empirically; Additional file 1: Fig. S14) and sketch size 10^7^. Finally, we use Equation 10 to compute the hamming distance with *η* and *ζ* values computed using Equations 6, 7 if *c* < 5 or else using Equation 9. The genome length *L* is estimated as the total sequence length divided by the coverage *c*.

### Experimental setup

#### Method settings

For Skmer, we use default parameters described above. For Mash, similar to Skmer, we used *k* = 31 (selected empirically; Additional file 1: Fig. S14) and sketch size 10^7^. As Mash handles errors by removing low copy *k*-mers, we set the minimum cardinality for *k*-mers to be included as 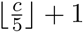 with our estimate of *c*.

AFF has an algorithm to correct hamming distances for low coverage, but the correction relies on adjusting the length of tip branches in a distance-based inferred phylogeny. As such, it cannot run on a pair of genomes and requires at least four genomes. Also, AAF leaves coverage estimation to the user with some guidelines, which we fully follow (Additional file 1: Appendix C).

For building phylogenetic distances, we we transformed Skmer distances using the JC69 [50] model and used FastME [49] to construct the distance-based trees via BIONJ [73] method.

#### Genomic Datasets

We used three sets of publicly available assembled genomes (Additional file 1: Tables S4–S6) and used ART [74] to simulate genome-skims of read length *ℓ* = 100 with default sequencing error profile, controlling for the sequencing depth (coverage) (Additional file 1: Appendix C). Specifically, the data included 21 *Drosophila* genomes (flies) and 22 genomes from the *Anopheles* genus (mosquitoes) obtained from InsectBase[75], and 47 avian species from the Avian Phylogenomic Project [76, 77].

For the experiment on real genome skims, high-coverage SRA’s of 14 *Drosophila* species were obtained from NCBI database under project number PRJNA427774 [78] and then subsampled to 100Mb. Assemblies used to compute true distances for these 14 *Drosophila* species were obtained from the Drosophila project [79]. We used the tool fastp [80] for filtering low-quality reads and adapter removal. We also used Megablast [81] to search against a database of bacterial and mitochondrial genomes and remove contaminant reads. We used Bowtie2 [63] with the highest sensitivity to remove the reads aligning (even partially) to the human reference genome.

To simulate genomes with controlled genomic distance, we introduced random mutations. As a challenging case, we took the highly repetitive assembly of the wasp species *Cotesia vestalis*, and mutated it artificially; we only applied single nucleotide mutations distributed uniformly at random across the genome. We repeated the study on the simpler case of the fly species *D. melanogaster*. We generate genome-skims using ART with *ℓ* = 100, default error profile of Illumina sequencer, and varying coverage between 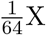 and 16X. For simulated genomes, we repeated the subsampling 10 times and reported the mean and standard error.

In order to compare with DNA barcoding method, we downloaded available COI barcodes for the *Drosophila* and *Anopheles* species in BOLD database [12]. Out of 21 *Drosophila* and 22 *Anopheles* species in our dataset, 16 *Drosophila* and 19 *Anopheles* species had one or more barcodes in BOLD. For each species, we selected a barcode, and using MUSCLE [82], aligned all barcodes within each dataset and constructed the phylogenetic tree assuming the Jukes-Cantor model. Under the same model of substitution, we transformed Skmer distances and built the Skmer tree. We used FastME [49] to construct the distance-based trees via BIONJ [73] method. The maximum-likelihood COI trees were built using PhyML [83].

#### Evaluation Metrics

For simulated data, the true distance is controlled and is thus known. For biological datasets, the ground truth is unknown. Instead, we use the distance measured on the full assembly by each method as its ground truth; thus, the ground truth for AAF is computed using AAF. We show both absolute error and the relative error, measured as 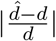 where *d* and 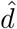 are the true and the estimated distances.

#### Leave-out

We used a leave-out strategy to study the accuracy of searching for a query genome in a reference set. For a query genome *G_q_* in a set of *n* genomes {*G*_1_… *G_n_*}, we ordered all genomes based on their distances to *G_q_* calculated using the full assemblies, which represents the ground truth; let 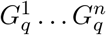 denote the order, and 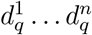 be the respective distances from the query (note 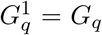 and 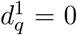). For 0.01 ≤ *d* ≤ 0.10, we removed genomes 1… *i* from the datasets where *i* is the largest value such that 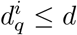, leaving us with 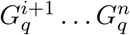. We then ordered the remaining genomes by each method; let *x*_1_… *x*_*n−i*_ be the order obtained by a method and let *r* be the the rank of the best remaining genome according to the ground truth in the estimated order (i.e., 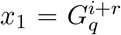). Since *r* = 1 implies perfect performance, and *r* > 1 indicates error, we measured rank error as the mean of *r* – 1 across all query genomes (1 ≤ *q* ≤ *n*). Moreover, the mean (relative) distance error is defined as the mean of 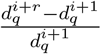 over all queries.

## Supplementary Material

### A Theoretical results

Consider two genomes of identical length *L* and separated by hamming distance *D* where the hamming distance is defined as the fraction of variant sites between the perfect alignment of the two genomes. We would like to estimate *D* from two genome-skims.

#### Mutations

We model the two genomes as the outcome of a random process that copies a genome and introduces mutations at each position i.i.d with a fixed probability *d*. Indexing from left to right, we can define *n* = *L* – *k* + 1 *k*-mers (note that *n* ≈ *L* for any reasonable choice of *k* and genome length). Let *X_i_* be a binary random variable (r.v.) that indicates whether *k*-mer *i* is identical between the two genomes. Clearly, in our model, *X_i_* ~ Bern(*p*) where *p* = (1 – *d*)^*k*^. Then, 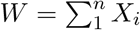 gives the number of shared *k*-mers. If *J* is defined as the Jaccard index over the set of all *k*-mers from both genomes, it’s easy to see that 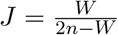 and thus, 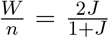. We further make a simplifying assumption. We assume all *X_i_* r.v.s are independent, an assumption that is true for most pairs of *k*-mers but ignores the fact that each *k*-mer overlaps with *k*-1 other *k*-mers. With this assumption, the maximum likelihood estimate of *p* is simply

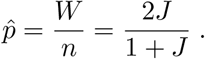

By the functional invariance of maximum likelihood, the ML estimate of *d* is given by:

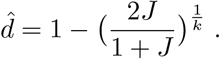

#### *k*-mer sampling

We now assume that each genome is covered uniformly at random. Thus, *k*-mers are also sub-sampled and we assume each *k*-mer is sampled at least once with probability *η*_1_ in the first genome and *η*_2_ in the second genome; we derive the relationship between these probabilities and genome coverage below. We estimate *η* values separately (also described below) and here consider them as given. For each 1 ≤ *i* ≤ *n* and *j* ∈ {1,2}, let *Y_j,i_* ~ Bern(*η_j_*) be the indicator of whether the *k*-mer *i* is sampled at least once in the genome *j*. Under this scenario, the number of *k*-mers shared between the two genomes is given by the r.v. 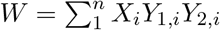. Defining *Z* = *X_i_Y*_1,*i*_*Y*_2,*i*_, we get 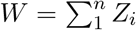 and *Z_i_* ~ Bern(*r*) where *r* = *pη*_1_*η*_2_ by the independence of the mutation process and each of the two *k*-mer sampling processes. Assuming independence between *Z_i_* r.v.s (again ignoring the overlap between consecutive *k*-mers) we get the ML estimate 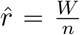, and thus (for a given *η*_1_ and *η*_2_) we have

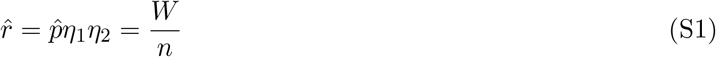

Let 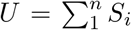 where *S_i_* = *Y*_1,*i*_ + *Y*_2,*i*_ – *Y*_1,*i*_*Y*_2,*i*_*X_i_*. It is easy to see that *U* gives the total number of sampled *k*-mers in both genomes. However, *S_i_* is not a Bernoulli and thus, *U* is not Binomial. Nevertheless, the same assumptions that we used to treat *X_i_* and *Z_i_* r.v.s as independent also give us independence between *S_i_* values; therefore, by the central limit theorem, 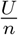 can be approximated by a Gaussian with mean 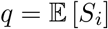. Moreover, 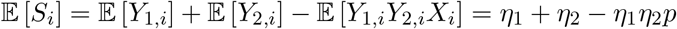 (note that *X_i_*, *Y*_1,*i*_ and *Y*_2,*i*_ are independent). By this Gaussian approximation, the ML estimate of *q* given *η*_1_,*η*_2_ is given by:

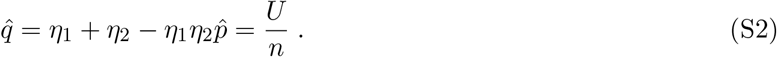

Note that 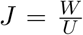. Equations S1 and S2 give two different ML estimators of the same parameter *p* given two different types of data (*W* and *U*). While the two estimators are not the same, because *n* is extremely large, both estimators have a very low variance. Exploiting the low variance, we treat the two estimates of *p* as equal and divide both sides of Equation S1 by Equation S2 to get:

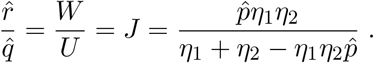

Solving for 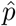 and replacing 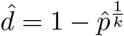 gives

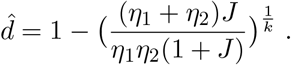

Note that we have assumed a known coverage and thus we are not co-estimating *η_j_*’s and *d*. In practice we need to first estimate *η*_1_ and *η*_2_, and we do it as we will describe.

#### Connection of *η* to read coverage

A *k*-mer stretching from position *y* to *y* + *k* on the genome is covered by the reads that start in the interval [*y* + *k* – *ℓ*, *y*]. Assuming that there is no sequencing error, and a uniform spread of of the *N* reads across the genome of length *L*. We show that the probability *η* that a *k*-mer is sampled by at least one read is give by

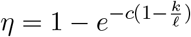

Let *X* be a r.v. denoting the number of reads that cover a specific *k*-mer. Assuming a uniform spread of *N* reads across the genome of length *L*, the probability of *x* reads covering a *k*-mer (starting in an interval of length *ℓ* – *k*) is given by

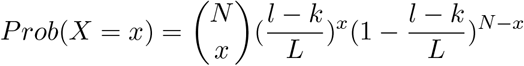

As *N* is large and 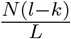 is constant, it can be closely approximated by

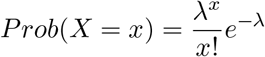

where 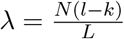 is the *k-mer* coverage, and is related to the coverage *c* by

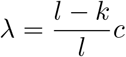

As the number of reads covering a *k*-mer follows Poisson distribution, the fraction of *k*-mers covered by 1 or more reads is

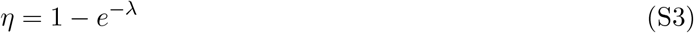

#### Sequencing error

We model the sequencing error as an i.i.d process that corrupts each position of each read with a fixed probability *ϵ*. To extend our previous results to cover this scenario, we need to see how the intersection r.v. (*W*) and the union r.v. (*U*) get affected.

We start with the intersection (*W*). We change the meaning of *η* to denote the probability that a *k*-mer is covered by at least one error-free read. The probability of a k-mer within a read being error-free is clearly

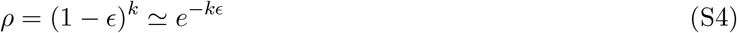

By conditioning on the number of reads covering a *k*-mer, the probability of not covering a *k*-mer with an error-free read is given by

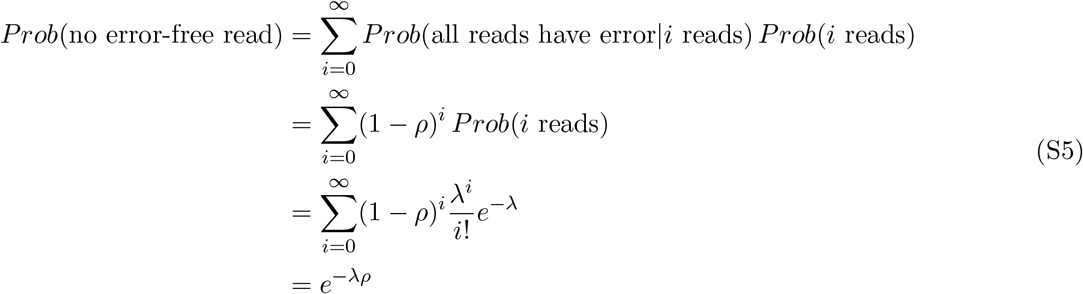

Hence, the probability that a *k*-mer is covered by at least one error-free read is given by

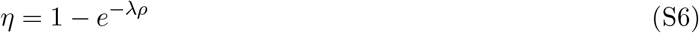

Note that Eqn. S6 reduces to Eqn. S3 when there is no sequencing error, i.e., *ρ* = 1. Similar to the case of no error, given *η*_1_ and *η*_2_, the r.v. 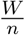 (where *W* is the number of shared *k*-mers) can be used with Equation S1 to estimate *r*.

We now turn to the union (r.v. *U*). For large enough *k*, and for genomes that are random and repeat-free, with high probability 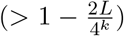 an error produces a new *k*-mer that is not observed in either of the input genomes. Ignoring the exceedingly unlikely event that two errors produce the same *k*-mer or that they produce a *k*-mer present in one of the two genomes, we can assume that the sequencing error generates as many new *k*-mers as the number of reads being affected by errors.

In the regime that includes errors, 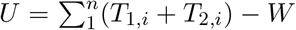 where the r.v.s *T*_1,*i*_ and *T*_2,*i*_ give the total number of *k*-mers generated from the position *i* from the first and second genomes, respectively. W.l.o.g, consider *T*_1,*i*_. By conditioning on the number of reads covering a *k*-mer we have

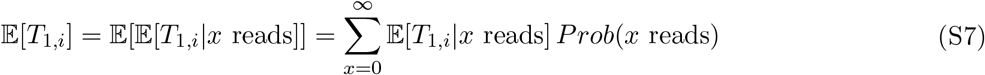

Given that *x* reads are covering a *k*-mer, *T*_1,*i*_ equals the number of erroneous *k*-mers *E*, plus 1 if there is any error-free *k*-mer. As *E* ~ *Binom*(*x*, 1 – *ρ*)

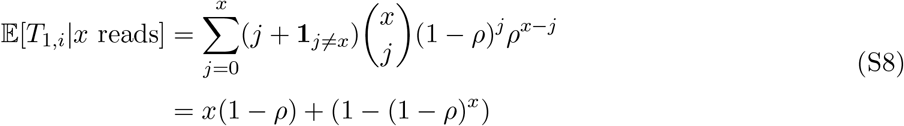

and substituting into (S7)

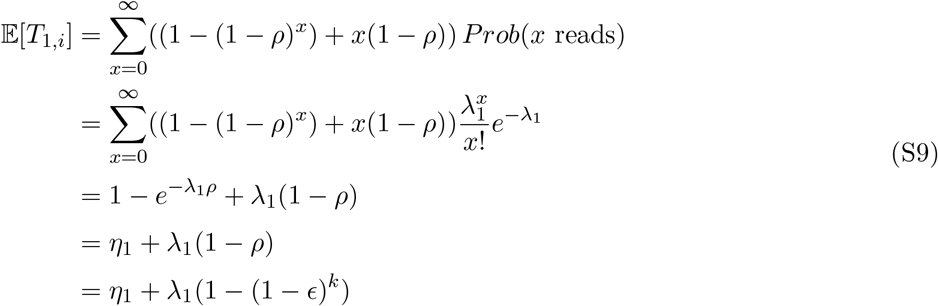

Letting 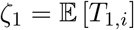 and using the same central limit argument we used before, 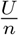 becomes approximately a Gaussian with expectation *ζ*_1_ + *ζ*_2_ – *η*_1_*η*_2_*p*. Similar to Equation S2, given *ζ*_1_, *ζ*_2_, *η*_1_, and *η*_2_, the Gaussian approximation gives us:

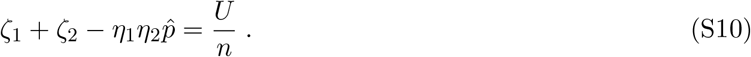

Again, assuming that estimates of *p* in Equation S1 (with the new definition of *η*) and Equation S10 are the same (due to low variance), we divide the two equations and solve for *d* to get the estimator:

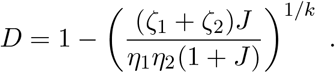

#### Excluding low-copy *k*-mers from the Jaccard index calculation

If we discard *k*-mers observed less than *m* times, then a *k*-mer will survive if it is covered by *m* or more error-free reads. Hence, *η* becomes the probability of *m* or more error-free reads covering a *k*-mer

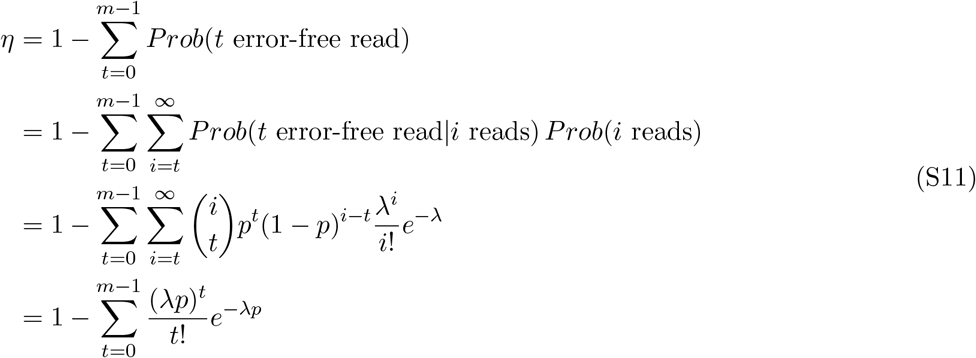

In general, we have shown that the probability distribution of the number of error-free k-mers is a Poisson with parameter λ*p*.

### B Computing GTR distances

To compute the GTR matrix using the log-det approach, we need a 4 × 4 matrix *F* where each element is the fraction of sites where one genome has one letter while the other genome has the other letter. Given this matrix, *d* = – log(det(*F*)).

As elsewhere, we assume a no-indel scenario so that each *k*-mer mismatch can be attributed to a single nucleotide substitution. For *i, j* ∈ {a,c,g,t}, let *x_ij_* = *x_ji_* denote the number of mutations of the form *i* ↔ *j*. Our goal is to estimate *x_ij_* for all *i, j*. However, the paradigm of computing distance by hashing/sketching *k*-mers treats all mutations alike. Formally, the estimated distance *d* equals

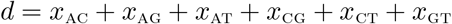

We do the following:

1. Replace *G* and *T* with *C*, and compute distance *d*_a_ = *x*_ac_ + *x*_ag_ + *x*_at_.
2. Replace *G* and *T* with *A*, and compute distance *d*_c_ = *x*_ac_ + *x*_cg_ + *x*_ct_.
3. Replace *G* with *T*, and compute distance *d*_ac_ = *x*_ac_ + *x*_ag_ + *x*_at_ + *x*_cg_ + *x*_ct_. Combining, we get

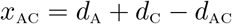 A similar procedure can be used to compute all *x_ij_* and normalization gives us *F*. Note that this procedure reduces the space of possible *k*-mers of length *k* to 2^*k*^ possibilities instead of 4^*k*^. Therefore, it will likely be required that *k* is increased for high accuracy when this approach is used.

### C Supplementary method details and commands

Here we provide the exact procedures and commands that we used to run external softwares throughout our experiments.

#### Simulating genome-skims using ART

To simulate short reads with length *ℓ* = 100 and (default) error profiles of Illumina HiSeq2000, we ran

~~~
art_illumina -i FASTA_FILE -l 100 -f c -o FASTQ_FILE
~~~

To simulate reads with constant error rate *ϵ* = 0.01 (Phred score = 20) at coverage *c*, we used

~~~
art_illumina -i FASTA_FILE -l 100 -qL 20 -qU 20 -f c -o FASTQ_FILE
~~~

#### Computing *k*-mer frequencies using JellyFish

To count all k-mers of length *k* = 31 in a genome-skim, we used

~~~
jellyfish count -m 31 -s 100M -C -o COUNT_FILE FASTQ_FILE
~~~

and to get the histogram of k-mer counts

~~~
jellyfish histo COUNT_FILE
~~~

#### Computing Jaccard index and estimating distance using Mash

We first *sketch* input genome-skims or assemblies with k-mer length *k* = 31 and sketch size *s* = 10^7^. For genome-skims (in FASTQ format) when no k-mer filtering is applied, we run

~~~
mash sketch -r -k 31 -s 10000000 -o SKETCH_FILE FASTQ_FILE
~~~

To sketch genome-skims while filtering k-mers with less than *C* copies, we use

~~~
mash sketch -m C -k 31 -s 10000000 -o SKETCH_FILE FASTQ_FILE
~~~

For genome assemblies (in FASTA format), we used

~~~
mash sketch -k 31 -s 10000000 -o SKETCH_FILE FASTA_FILE
~~~

Then, the Jaccard index and Mash distance between sketches is computed by running

~~~
mash dist SKETCH_FILE_1 SKETCH_FILE_2
~~~

#### Estimating distances using AAF

To count the k-mers (*k* = 31) in a dataset of genome-skims using 24 cores and 120GB memory, we first ran

~~~
python PATH_to_FILE/aaf_phylokmer.py -k 31 -t 24 -o KMER_COUNT_FILE \ -d INPUT_DIR -G 120
~~~

Next, to get the (uncorrected) distances and phylogeny, we used

~~~
python PATH_to_FILE/aaf_distance.py -i KMER_COUNT_FILE -t 24 -G 120 \ -o OUTPUT_FILE_PREFIX -f KMER_DIVERSITY_FILE
~~~

where KMER_DIVERSITY_FILE is an output of previous command. Finally, to correct tip branches of phylogeny tree for low coverage and sequencing error, we used

~~~
python PATH_to_FILE/aaf_tip.py -i TREE_FILE -k 31 \ --tip TIP_INFO_FILE -f KMER_DIVERSITY_FILE
~~~

where we had to provide `TIP_INFO_FILE` containing estimates of coverage and sequencing error. To estimate coverage, we followed the procedure suggested in AAF user manual. We first used JellyFish to find the k-mer counts *M_i_*’s as described before. They suggest when there is a clear peak in the *k*-mer frequency distribution, estimate *k*-mer coverage λ to be the maximum bin. As they do not suggest a specific rule for that, we first find *j* = argmax_*i*>1_ *M_i_*, excluding the count of the first bin *M*_1_, which is always large because of erroneous k-mers due to sequencing error. If *j* > 2, it means that we can see a peak in k-mers distribution at *j*, so we use λ = *j*. Otherwise, if *j* = 2, we follow their suggested formula 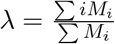 for the case of low coverage or high sequencing error that there is no clear peak in the k-mer frequency distribution. We should also mention that no k-mer filtering used for AAF, as the coverage was heterogeneous over genome-skims. In fact, in AAF the filtering is applied to all genome-skims if used, and so they suggest to not apply filtering when there is any taxon with low coverage (*c* < 5) within the dataset.

**Figure S1:**
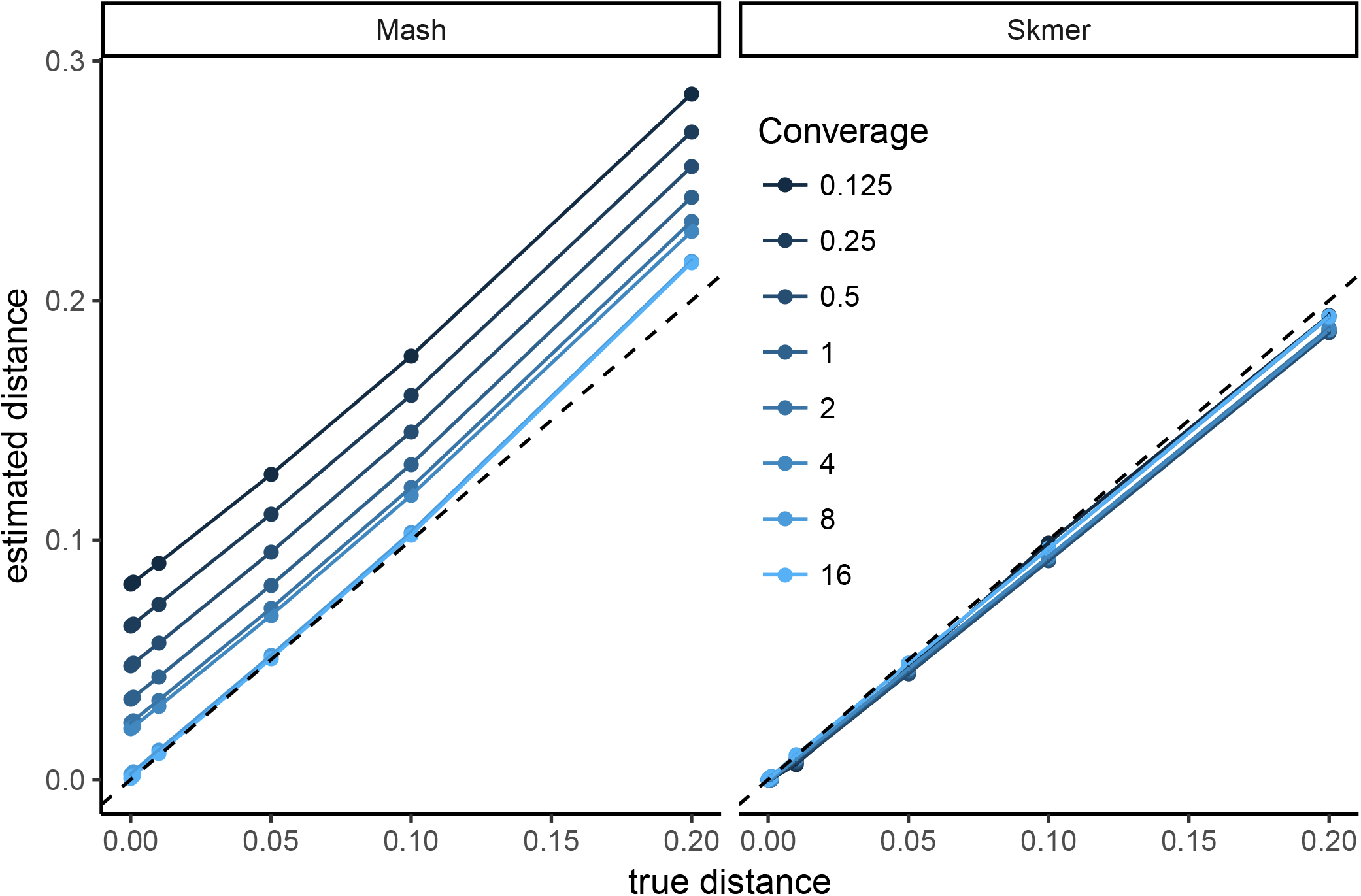
Comparing the accuracy of Mash and Skmer on simulated genomes. Genome-skims are simulated using ART with read length *ℓ* = 100. Substitutions applied to the assembly of *C. vestalis* at six different rates (x-axis), and genome-skims simulated at varying coverage range from 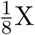 to 16X (colors). The estimated distance (y-axis) by Mash (left) and Skmer (right) is plotted versus the real distances (x-axis). The mean (dots) distances are shown as dots (10 repeats) but standard errors are too small to see. The unit line is shown as a dashed line.

**Figure S2:**
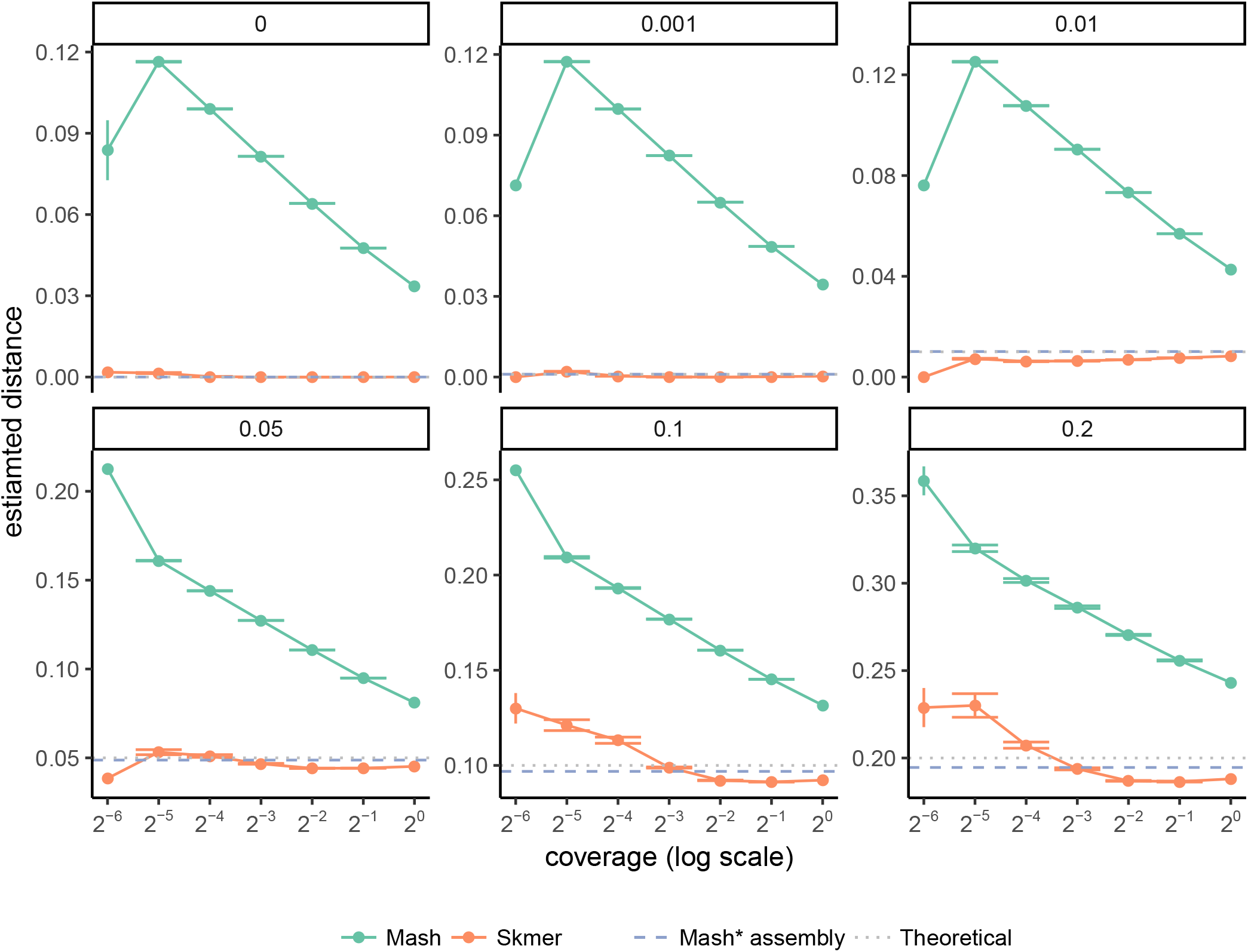
Comparing distances estimated by Mash and Skmer for simulated data at very low coverages. Skims of *C. vestalis* v.s. genomes simulated to be at different distances from *C. vestalis*, with varying coverage. The mean and standard error of distances are shown over 10 repeats of the experiment. The coverage ranges from 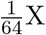 to 1X.

**Figure S3:**
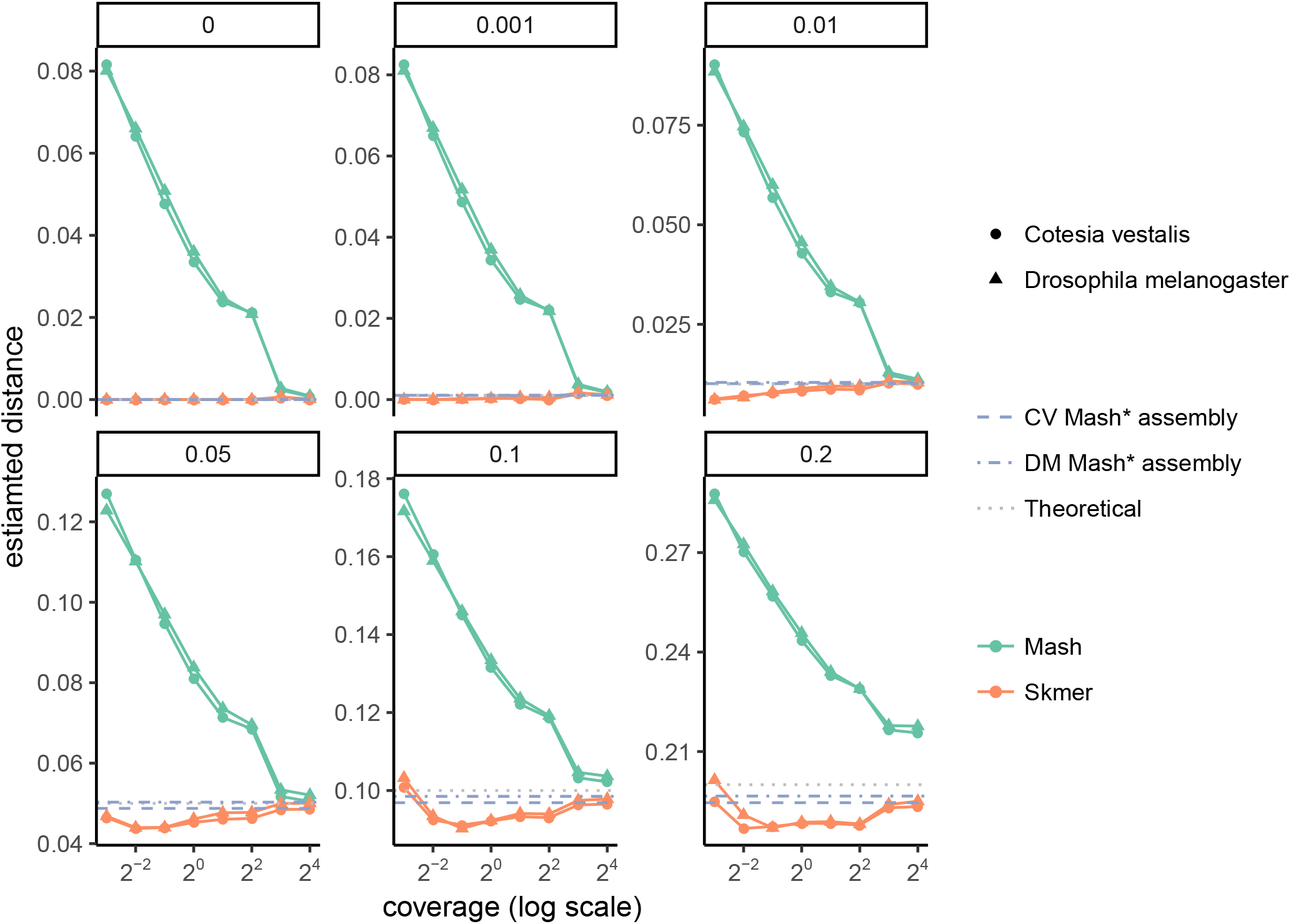
Comparing distances estimated for genome-skims of two different species. Genomes simulated at different distances from the genomes of C. vestalis and D. melanogaster and subsampled at a range of coverage from 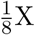 to 16X.

**Figure S4:**
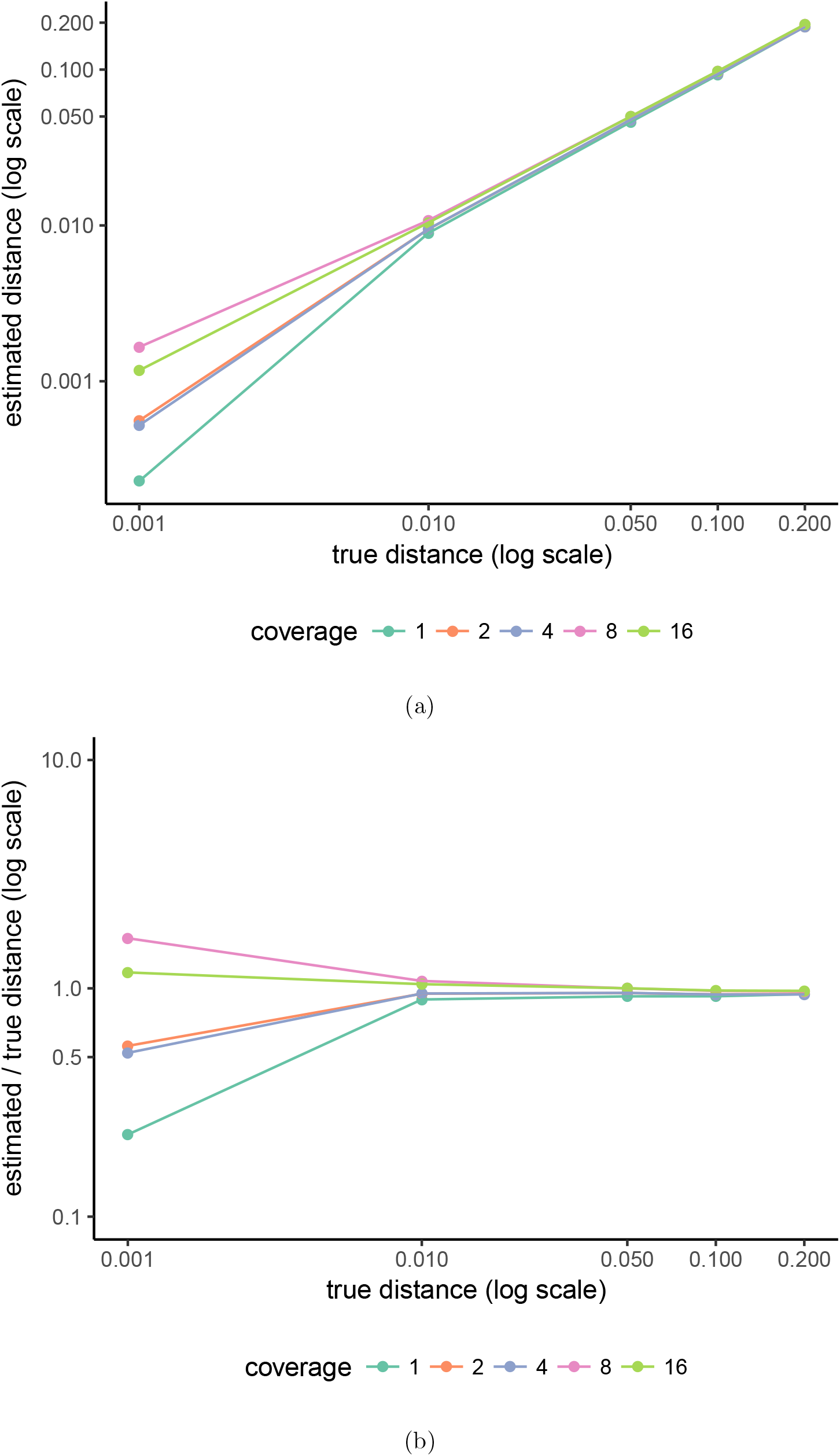
The resolution of Skmer at different genomic distances. Skims of *D. melanogaster* v.s. genomes simulated to be at different distances from *D. melanogaster*, with varying coverage. (a) Estimated distance versus the true distance. (b) The ratio of estimated distance to the true distance.

**Figure S5:**
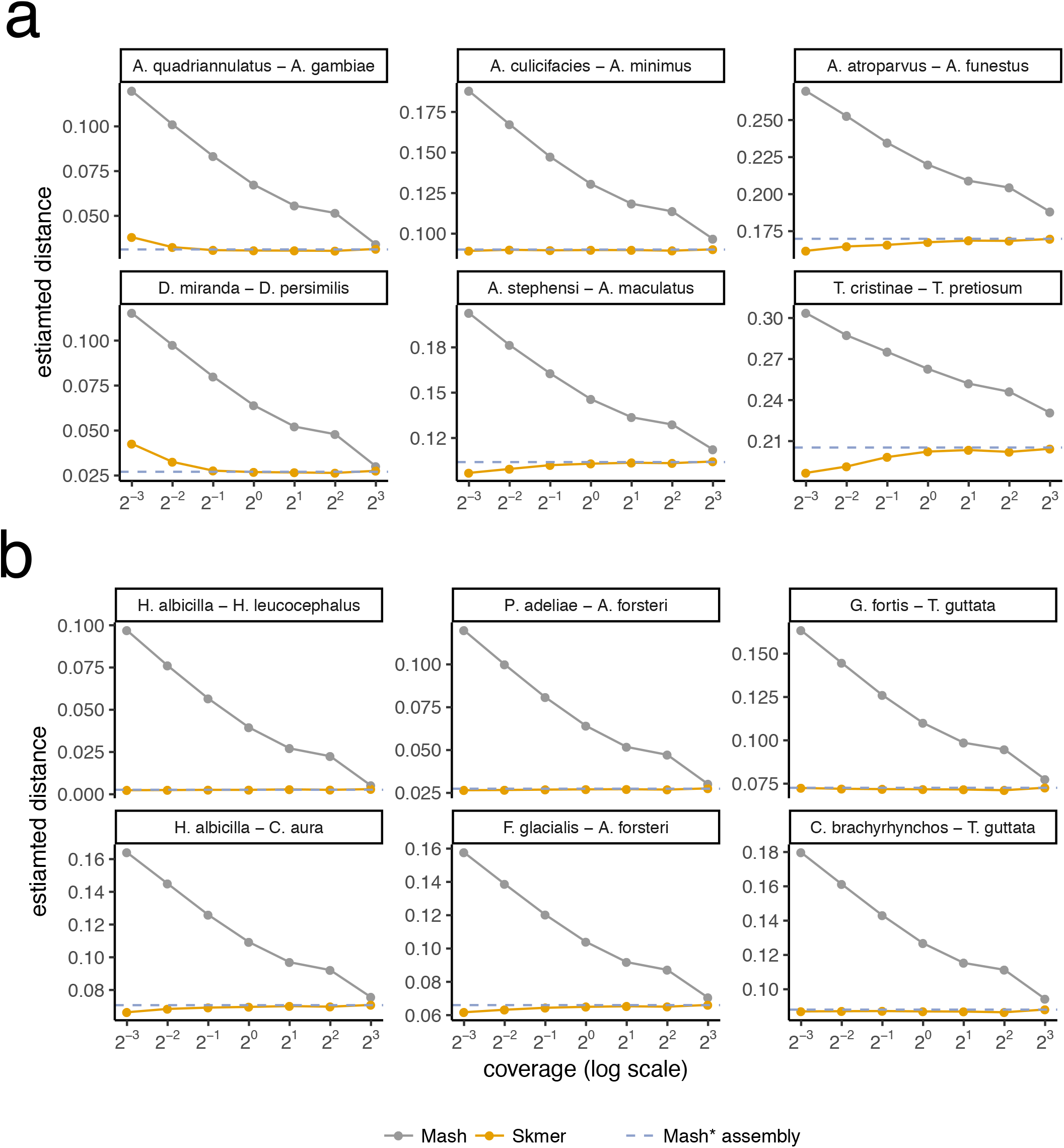
Comparing the accuracy of Mash and Skmer on pairs of insects and birds genomes. Genome-skims simulated at coverage 8X to 8X. On each subplot, the estimated distance (y-axis) is plotted versus the coverage (x-axis) for a pair of species. Dashed line shows Mash* run on assemblies, which is taken as the true distance. Skmer estimates (light-colored curves) are very close to the true distance while Mash (gray curves) largely overestimates at lower coverages. (a) Six pairs of insects. (b) Six pairs of birds.

**Figure S6:**
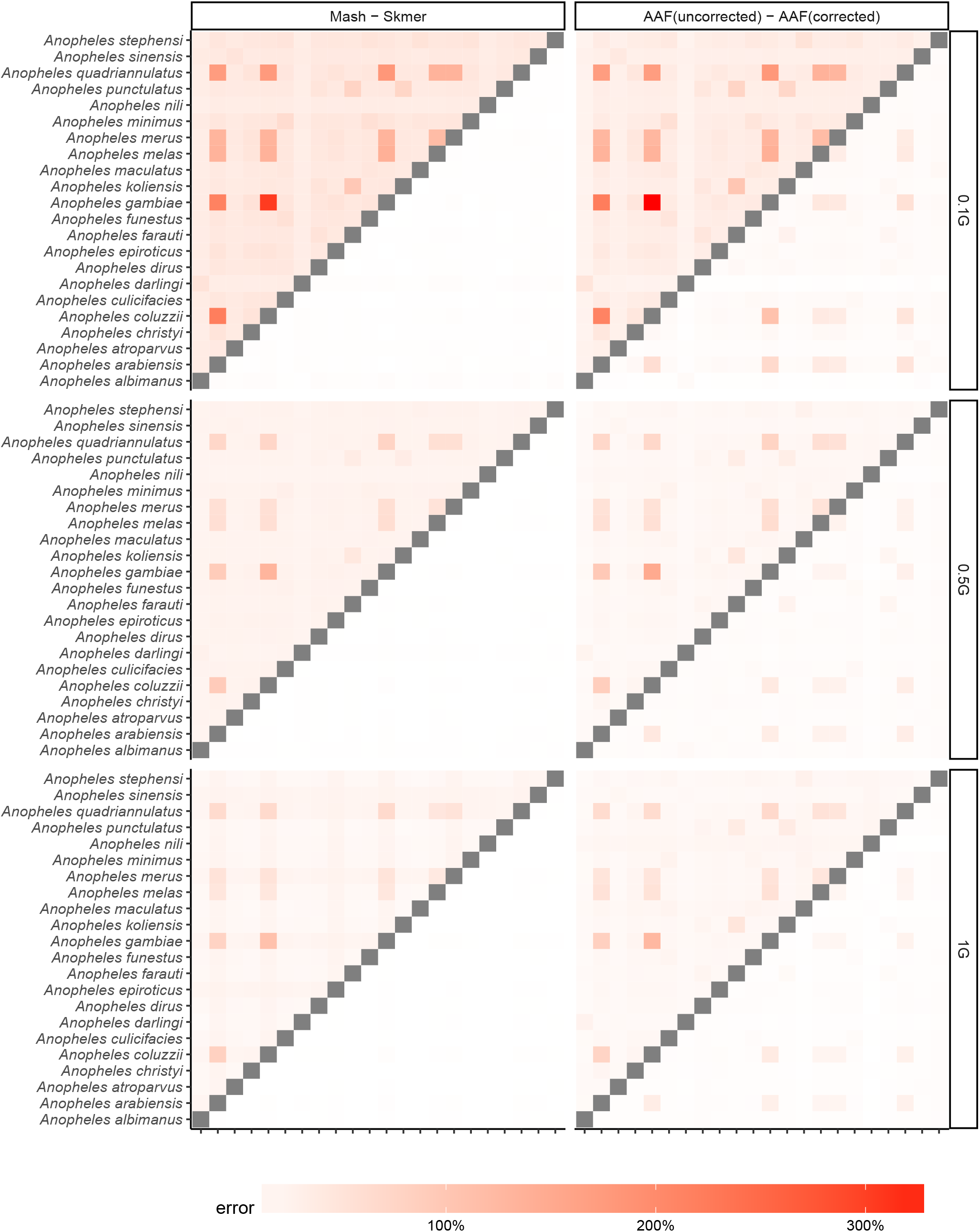
Comparing the error of Mash, Skmer, and AAF in distance estimation with fixed amount of sequence from each species. The dataset of 22 Anopheles genomes, subsampled with 0.1Gb, 0.5Gb, and 1Gb sequence.

**Figure S7:**
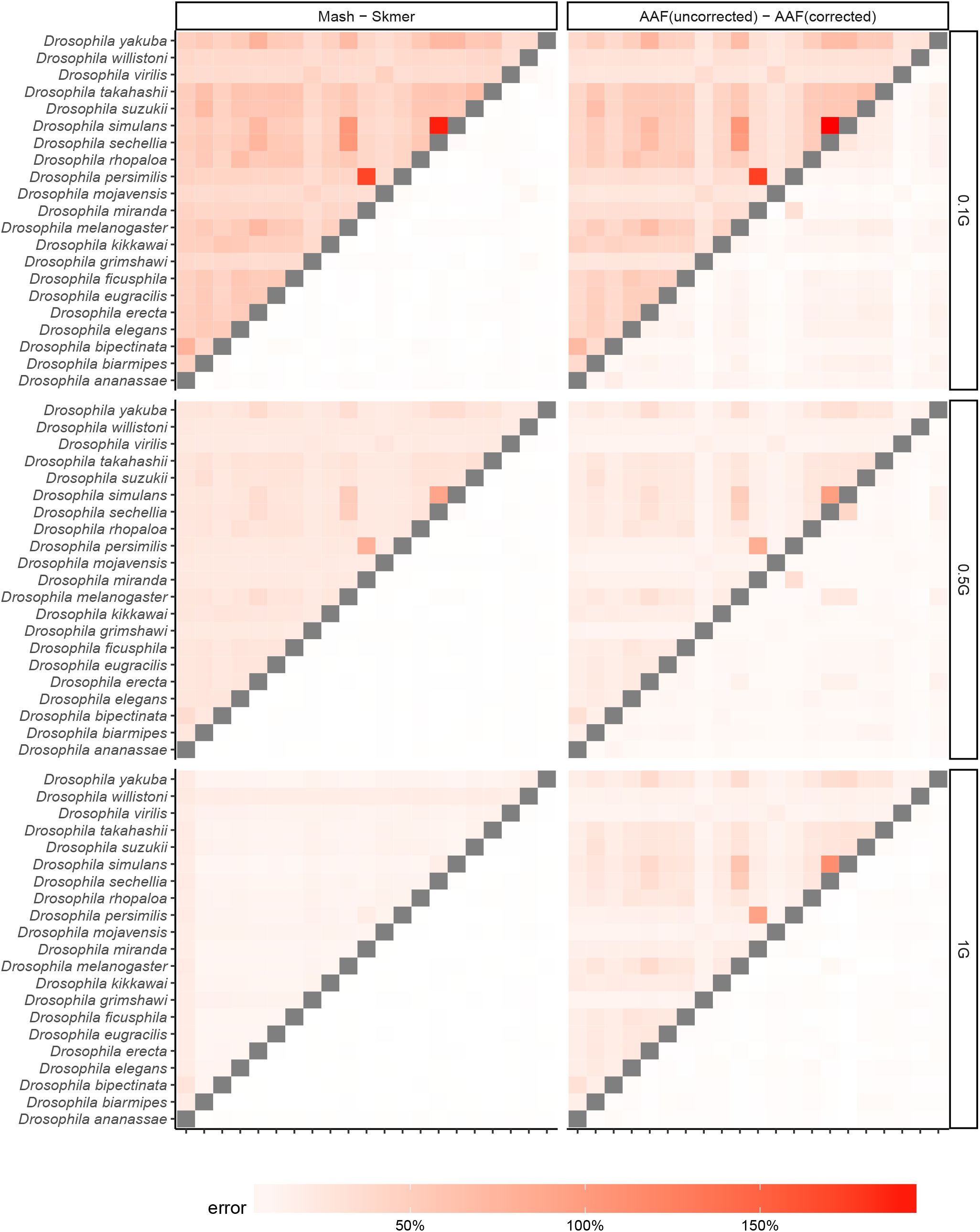
Comparing the error of Mash, Skmer, and AAF in distance estimation with fixed amount of sequence from each species. The dataset of 21 Drosophila genomes, subsampled with 0.1Gb, 0.5Gb, and 1Gb sequence.

**Figure S8:**
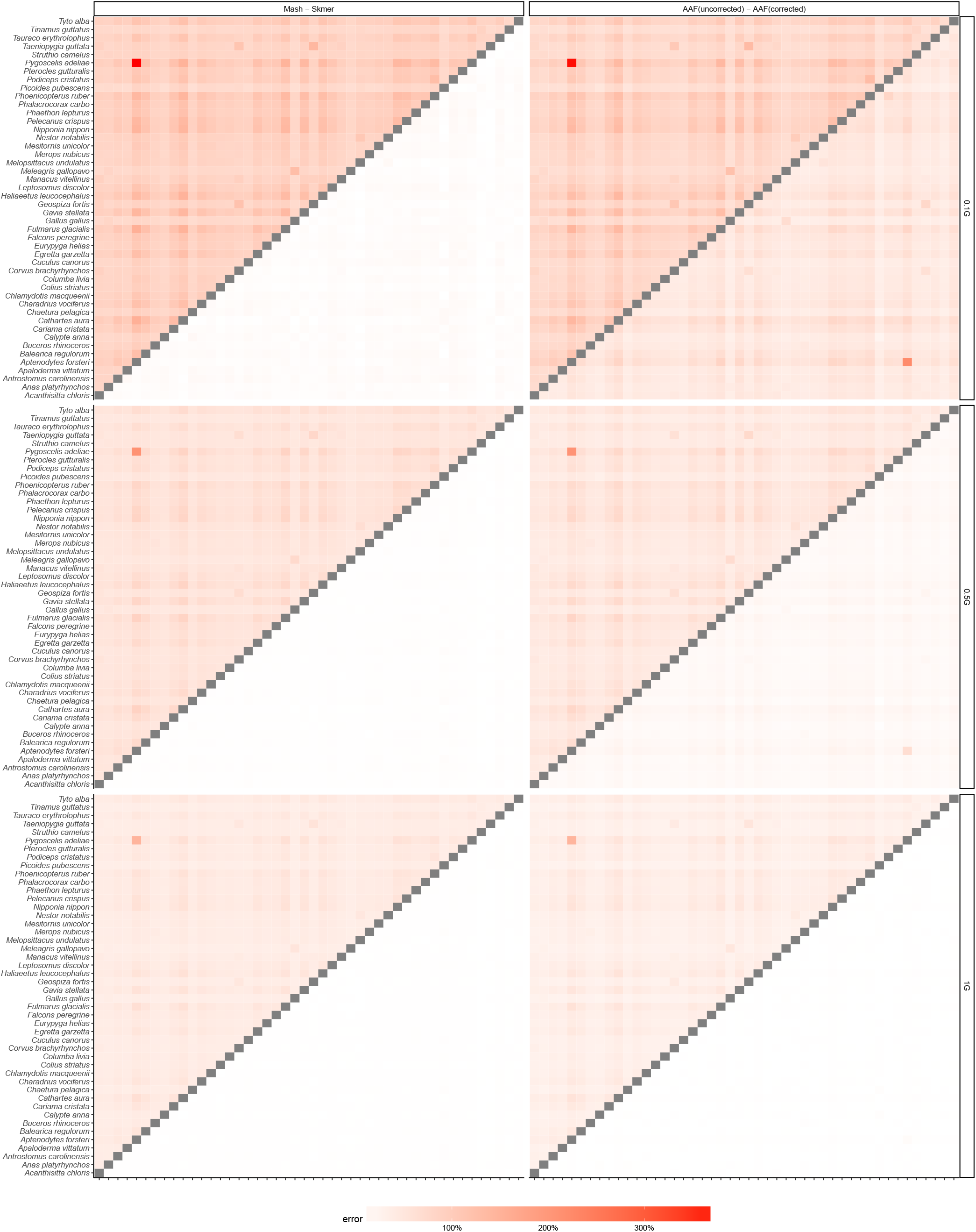
Comparing the error of Mash, Skmer, and AAF in distance estimation with fixed amount of sequence from each species. The dataset of 47 avian genomes, subsampled with 0.1Gb, 0.5Gb, and 1Gb sequence.

**Figure S9:**
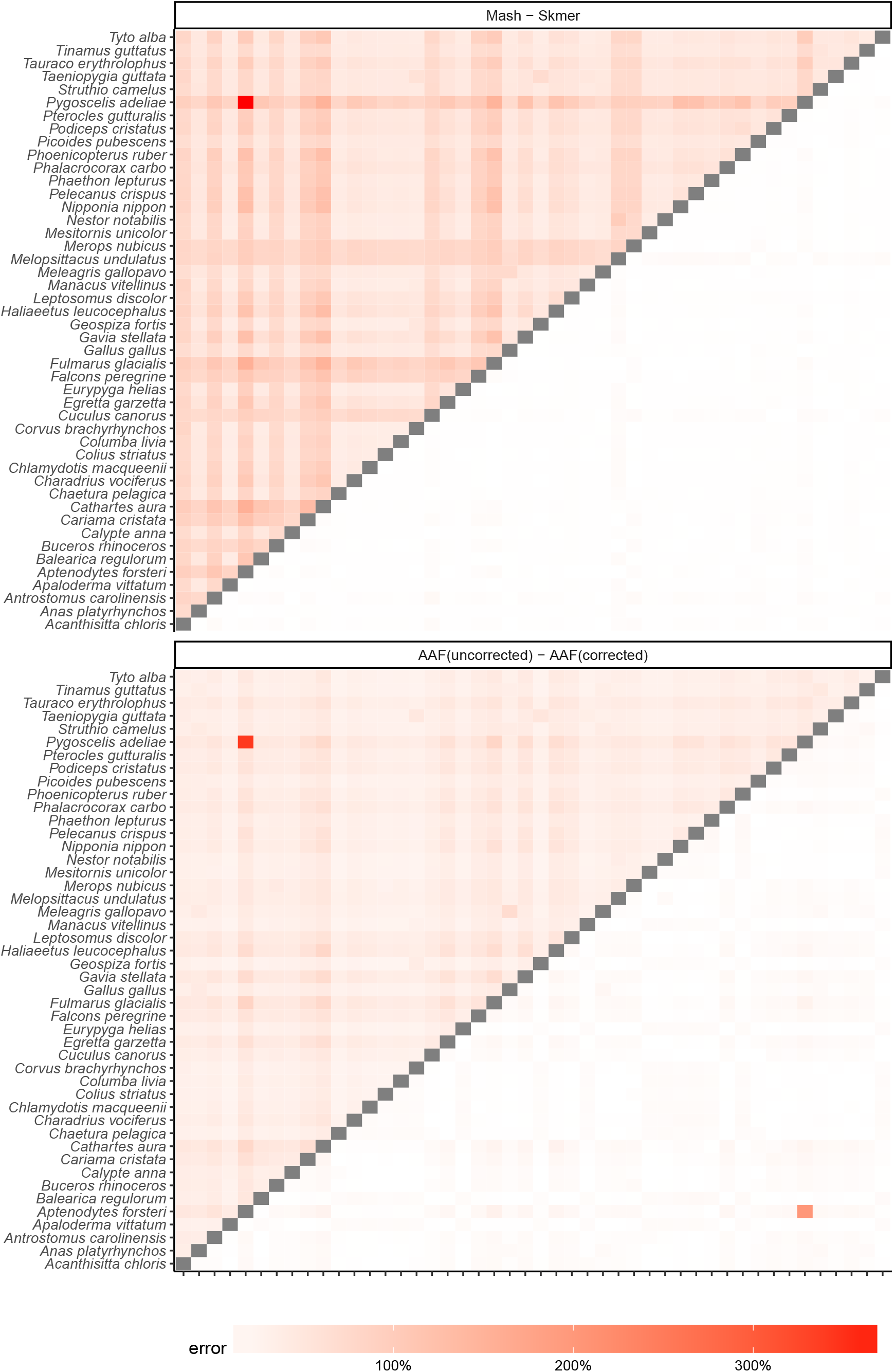
Comparing the error of Mash, Skmer, and AAF on the Avian dataset with mixed coverage. Species have random amount of sequence chosen uniformly among 0.1Gb, 0.5Gb, and 1Gb. Similar to (Fig. 5), we have excluded one of the eagles (*H. albicilla*). The error of Mash, AAF, and Skmer in estimating the distance between the two eagles are 2193%, 884%, and 4.2%, respectively (both of the eagles are subsampled at 0.5Gb here).

**Figure S10:**
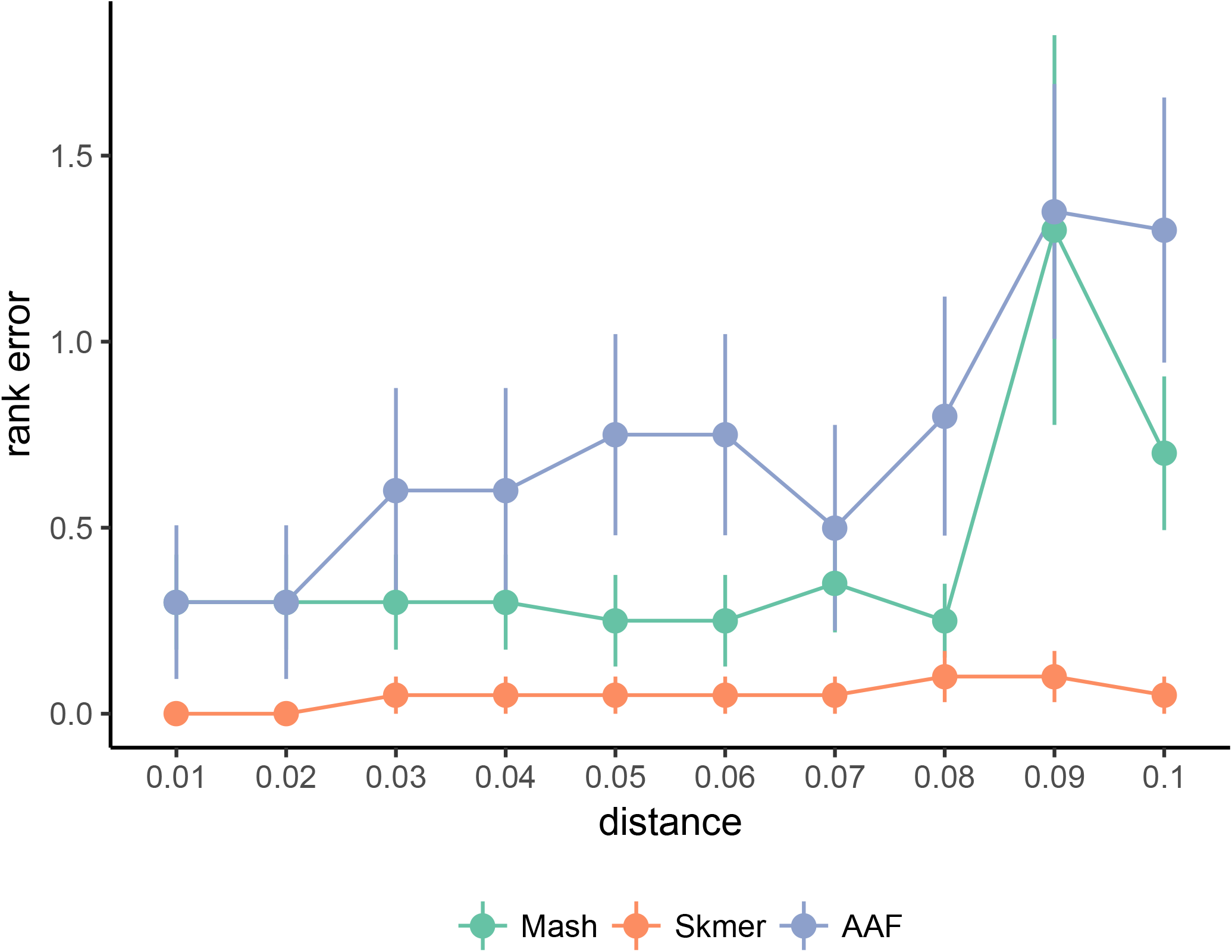
The mean rank error of the best remaining match in leave-out experiments on the *Drosophila* dataset. *Drosophila willistoni* has been excluded.

**Figure S11:**
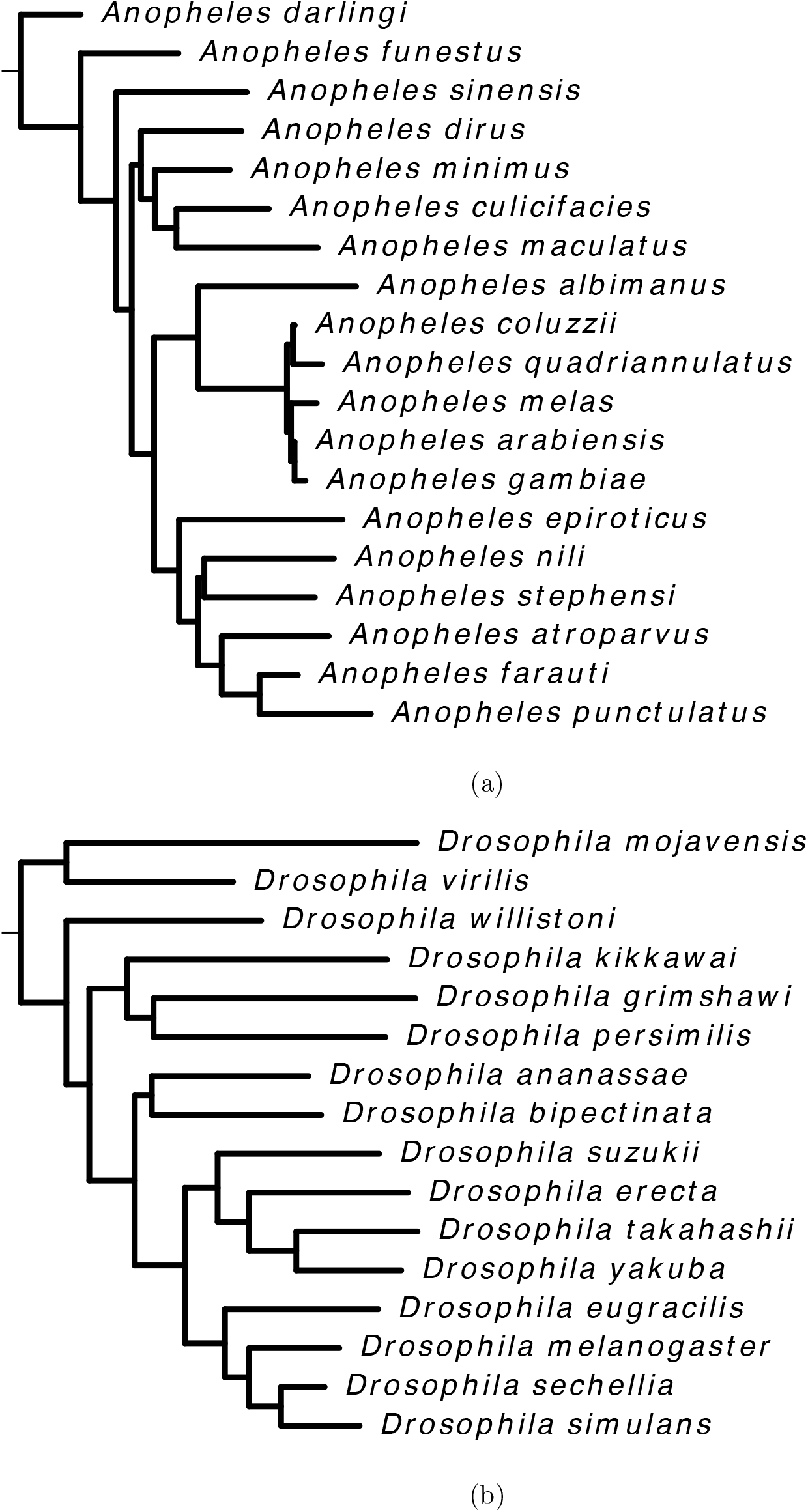
Maximum-likelihood trees inferred from COI barcodes. (a) *Anopheles* tree. (b) *Drosophila* tree.

**Figure S12:**
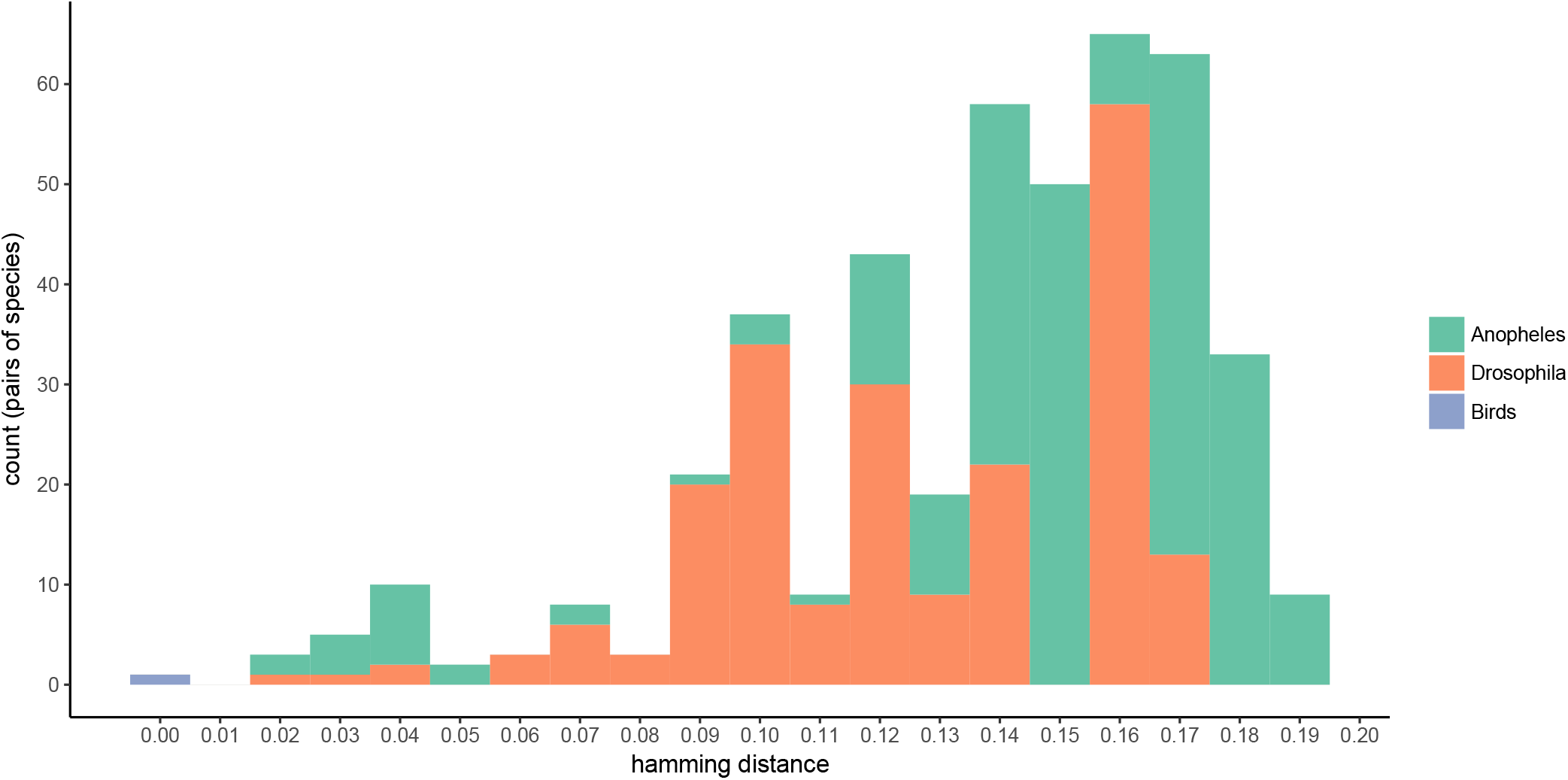
The histogram of genomic distances between species from the same genus among the Anopheles, Drosophila, and birds datasets. Distances computed based on full assemblies. The only species from the same genus with hamming distance less than 0.01 were the two eagle species *(H. albicilla* and *H. leucocephalus*).

**Figure S13:**
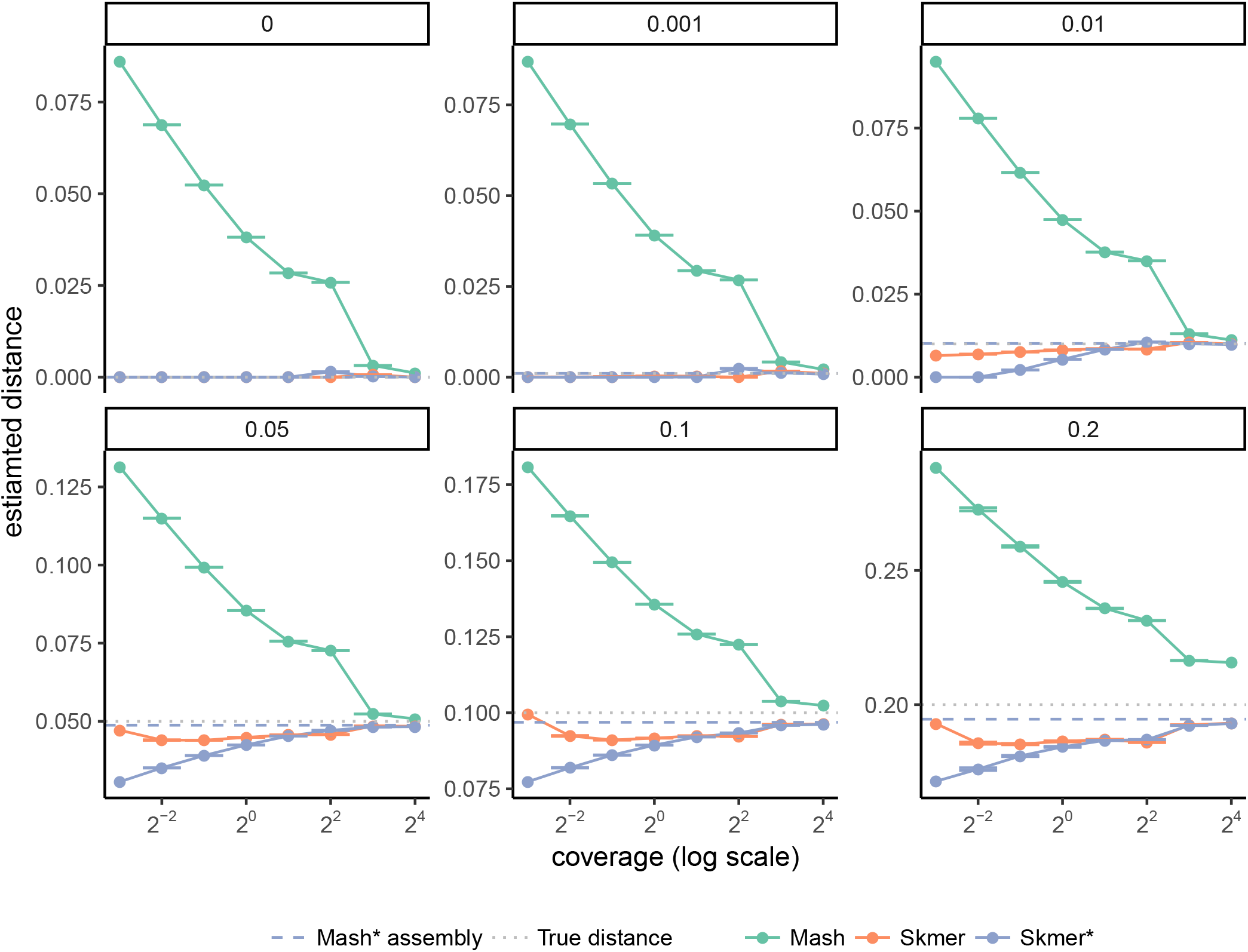
The performance of Skmer coverage estimation. Comparing distances estimated by Mash, Skmer with estimated coverages, and Skmer with true coverages (Skmer*), on genome-skims of *C. vestalis* and genomes simulated at different distances from it.

**Figure S14:**
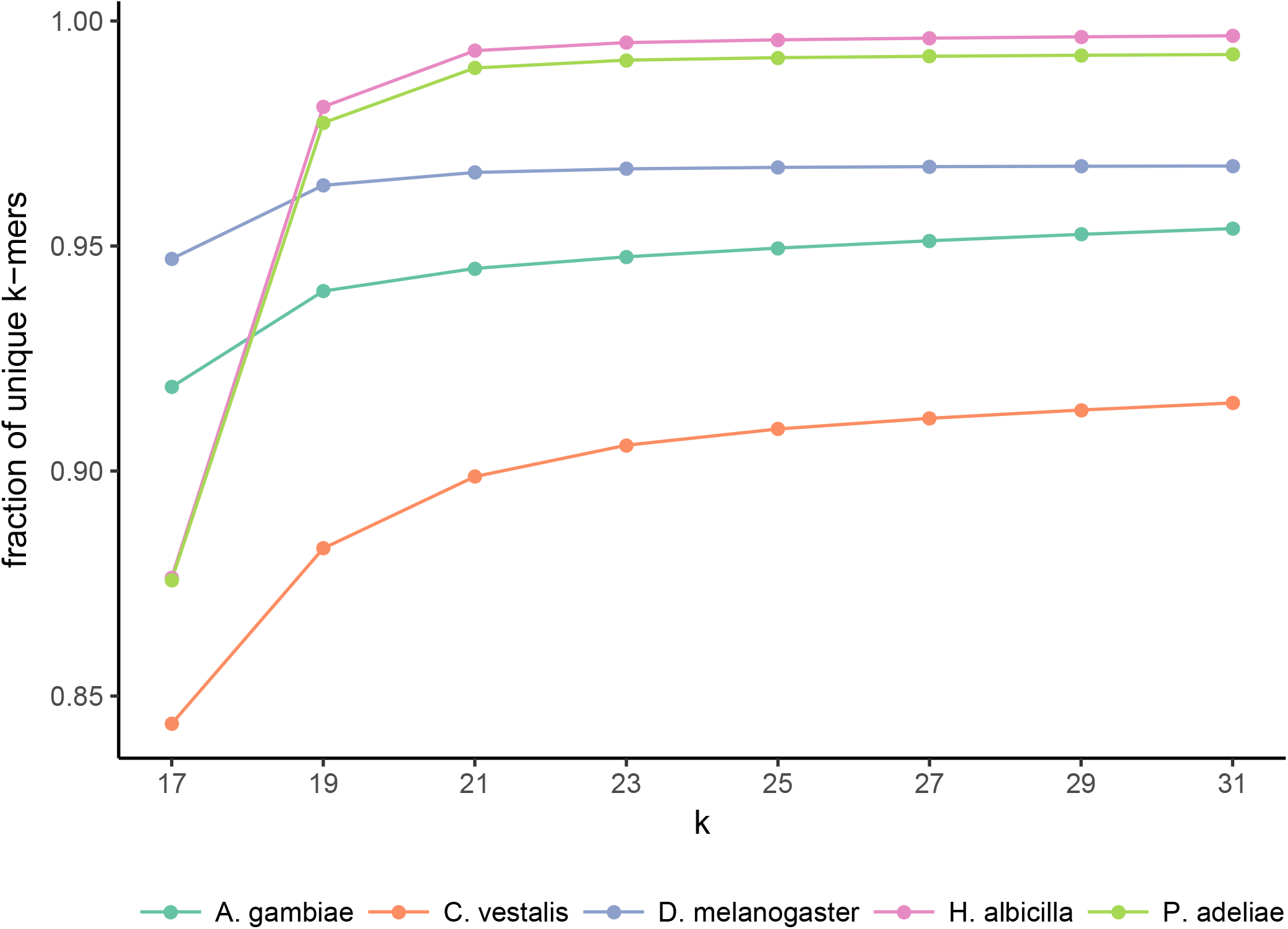
The fraction of unique *k*-mers in selected species of insects and birds.

**Table S3:**
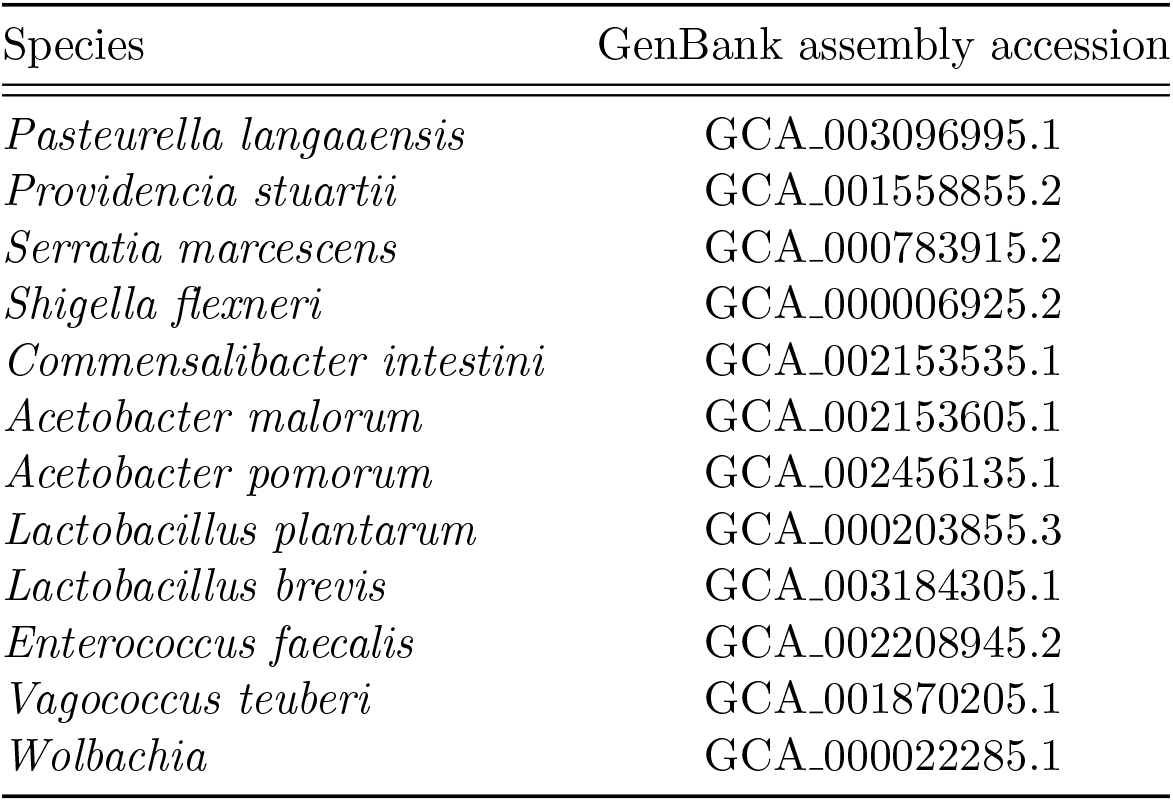
GenBank accession numbers of microbial species used in contamination removal.

**Table S4:** GenBank accession numbers and URLs for Anopheles genomes

| Species | GenBank assembly accession | URL |
| --- | --- | --- |
| *Anopheles albimanus* | GCA_000349125.1 | http://www.insect-genome.com/data/genome_download/Anopheles_albimanus/Anopheles_albimanus_genomic.fasta.gz |
| *Anopheles arabiensis* | GCA_000349185.1 | http://www.insect-genome.com/data/genome_download/Anopheles_arabiensis/Anopheles_arabiensis_genomic.fasta.gz |
| *Anopheles atroparvus* | GCA_000473505.1 | http://www.insect-genome.com/data/genome_download/Anopheles_atroparvus/Anopheles_atroparvus_genomic.fasta.gz |
| *Anopheles christyi* | GCA_000349165.1 | http://www.insect-genome.com/data/genome_download/Anopheles_christyi/Anopheles_christyi_genomic.fasta.gz |
| *Anopheles coluzzii* | - | http://www.insect-genome.com/data/genome_download/Anopheles_coluzzii/Anopheles_coluzzii_genomic.fasta.gz |
| *Anopheles culicifacies* | GCA_000473375.1 | http://www.insect-genome.com/data/genome_download/Anopheles_culicifacies/Anopheles_culicifacies_genomic.fasta.gz |
| *Anopheles darlingi* | GCA_000211455.3 | http://www.insect-genome.com/data/genome_download/Anopheles_darlingi/Anopheles_darlingi_genomic.fasta.gz |
| *Anopheles dirus* | GCA_000349145.1 | http://www.insect-genome.com/data/genome_download/Anopheles_dirus/Anopheles_dirus_genomic.fasta.gz |
| *Anopheles epiroticus* | GCA_000349105.1 | http://www.insect-genome.com/data/genome_download/Anopheles_epiroticus/Anopheles_epiroticus_genomic.fasta.gz |
| *Anopheles farauti* | GCA_000956265.1 | http://www.insect-genome.com/data/genome_download/Anopheles_farauti/Anopheles_farauti_genomic.fasta.gz |
| *Anopheles funestus* | GCA_000349085.1 | http://www.insect-genome.com/data/genome_download/Anopheles_funestus/Anopheles_funestus_genomic.fasta.gz |
| *Anopheles gambiae* | GCA_000150785.1 | http://www.insect-genome.com/data/genome_download/Anopheles_gambiae/Anopheles_gambiae_genomic.fasta.gz |
| *Anopheles koliensis* | GCA_000956275.1 | http://www.insect-genome.com/data/genome_download/Anopheles_koliensis/Anopheles_koliensis_genomic.fasta.gz |
| *Anopheles maculatus* | GCA_000473185.1 | http://www.insect-genome.com/data/genome_download/Anopheles_maculatus/Anopheles_maculatus_genomic.fasta.gz |
| *Anopheles melas* | GCA_000473525.2 | http://www.insect-genome.com/data/genome_download/Anopheles_melas/Anopheles_melas_genomic.fasta.gz |
| *Anopheles merus* | GCA_000473845.2 | http://www.insect-genome.com/data/genome_download/Anopheles_merus/Anopheles_merus_genomic.fasta.gz |
| *Anopheles minimus* | GCA_000349025.1 | http://www.insect-genome.com/data/genome_download/Anopheles_minimus/Anopheles_minimus_genomic.fasta.gz |
| *Anopheles nili* | GCA_000439205.1 | http://www.insect-genome.com/data/genome_download/Anopheles_nili/Anopheles_nili_genomic.fasta.gz |
| *Anopheles punctulatus* | GCA_000956255.1 | http://www.insect-genome.com/data/genome_download/Anopheles_punctulatus/Anopheles_punctulatus_genomic.fasta.gz |
| *Anopheles quadriannulatus* | GCA_000349065.1 | http://www.insect-genome.com/data/genome_download/Anopheles_quadriannulatus/Anopheles_quadriannulatus_genomic.fasta.gz |
| *Anopheles sinensis* | GCA_000441895.2 | http://www.insect-genome.com/data/genome_download/Anopheles_sinensis/Anopheles_sinensis_genomic.fasta.gz |
| *Anopheles stephensi* | GCA_000300775.2 | http://www.insect-genome.com/data/genome_download/Anopheles_stephensi/Anopheles_stephensi_genomic.fasta.gz |

**Table S5:** GenBank accession numbers and URLs for Drosophila genomes

| Species | GenBank assembly accession | URL |
| --- | --- | --- |
| *Drosophila ananassae* | GCA_000005115.1 | http://www.insect-genome.com/data/genome_download/Drosophila_ananassae/Drosophila_ananassae_genomic.fasta.gz |
| *Drosophila biarmipes* | GCA_000233415.2 | http://www.insect-genome.com/data/genome_download/Drosophila_biarmipes/Drosophila_biarmipes_genomic.fasta.gz |
| *Drosophila bipectinata* | GCA_000236285.2 | http://www.insect-genome.com/data/genome_download/Drosophila_bipectinata/Drosophila_bipectinata_genomic.fasta.gz |
| *Drosophila elegans* | GCA_000224195.2 | http://www.insect-genome.com/data/genome_download/Drosophila_elegans/Drosophila_elegans_genomic.fasta.gz |
| *Drosophila erecta* | GCA_000005135.1 | http://www.insect-genome.com/data/genome_download/Drosophila_erecta/Drosophila_erecta_genomic.fasta.gz |
| *Drosophila eugracilis* | GCA_000236325.2 | http://www.insect-genome.com/data/genome_download/Drosophila_eugracilis/Drosophila_eugracilis_genomic.fasta.gz |
| *Drosophila ficusphila* | GCA_000220665.2 | http://www.insect-genome.com/data/genome_download/Drosophila_ficusphila/Drosophila_ficusphila_genomic.fasta.gz |
| *Drosophila grimshawi* | GCA_000005155.1 | http://www.insect-genome.com/data/genome_download/Drosophila_grimshawi/Drosophila_grimshawi_genomic.fasta.gz |
| *Drosophila kikkawai* | GCA_000224215.2 | http://www.insect-genome.com/data/genome_download/Drosophila_kikkawai/Drosophila_kikkawai_genomic.fasta.gz |
| *Drosophila melanogaster* | GCA_000778455.1 | http://www.insect-genome.com/data/genome_download/Drosophila_melanogaster/Drosophila_melanogaster_genomic.fasta.gz |
| *Drosophila miranda* | GCA_000269505.2 | http://www.insect-genome.com/data/genome_download/Drosophila_miranda/Drosophila_miranda_genomic.fasta.gz |
| *Drosophila mojavensis* | GCA_000005175.1 | http://www.insect-genome.com/data/genome_download/Drosophila_mojavensis/Drosophila_mojavensis_genomic.fasta.gz |
| *Drosophila persimilis* | GCA_000005195.1 | http://www.insect-genome.com/data/genome_download/Drosophila_persimilis/Drosophila_persimilis_genomic.fasta.gz |
| *Drosophila rhopaloa* | GCA_000236305.2 | http://www.insect-genome.com/data/genome_download/Drosophila_rhopaloa/Drosophila_rhopaloa_genomic.fasta.gz |
| *Drosophila sechellia* | GCA_000005215.1 | http://www.insect-genome.com/data/genome_download/Drosophila_sechellia/Drosophila_sechellia_genomic.fasta.gz |
| *Drosophila simulans* | GCA_000259055.1 | http://www.insect-genome.com/data/genome_download/Drosophila_simulans/Drosophila_simulans_genomic.fasta.gz |
| *Drosophila suzukii* | GCA_000472105.1 | http://www.insect-genome.com/data/genome_download/Drosophila_suzukii/Drosophila_suzukii_genomic.fasta.gz |
| *Drosophila takahashii* | GCA_000224235.2 | http://www.insect-genome.com/data/genome_download/Drosophila_takahashii/Drosophila_takahashii_genomic.fasta.gz |
| *Drosophila virilis* | GCA_000005245.1 | http://www.insect-genome.com/data/genome_download/Drosophila_virilis/Drosophila_virilis_genomic.fasta.gz |
| *Drosophila willistoni* | GCA_000005925.1 | http://www.insect-genome.com/data/genome_download/Drosophila_willistoni/Drosophila_willistoni_genomic.fasta.gz |
| *Drosophila yakuba* | GCA_000005975.1 | http://www.insect-genome.com/data/genome_download/Drosophila_yakuba/Drosophila_yakuba_genomic.fasta.gz |

**Table S6:**
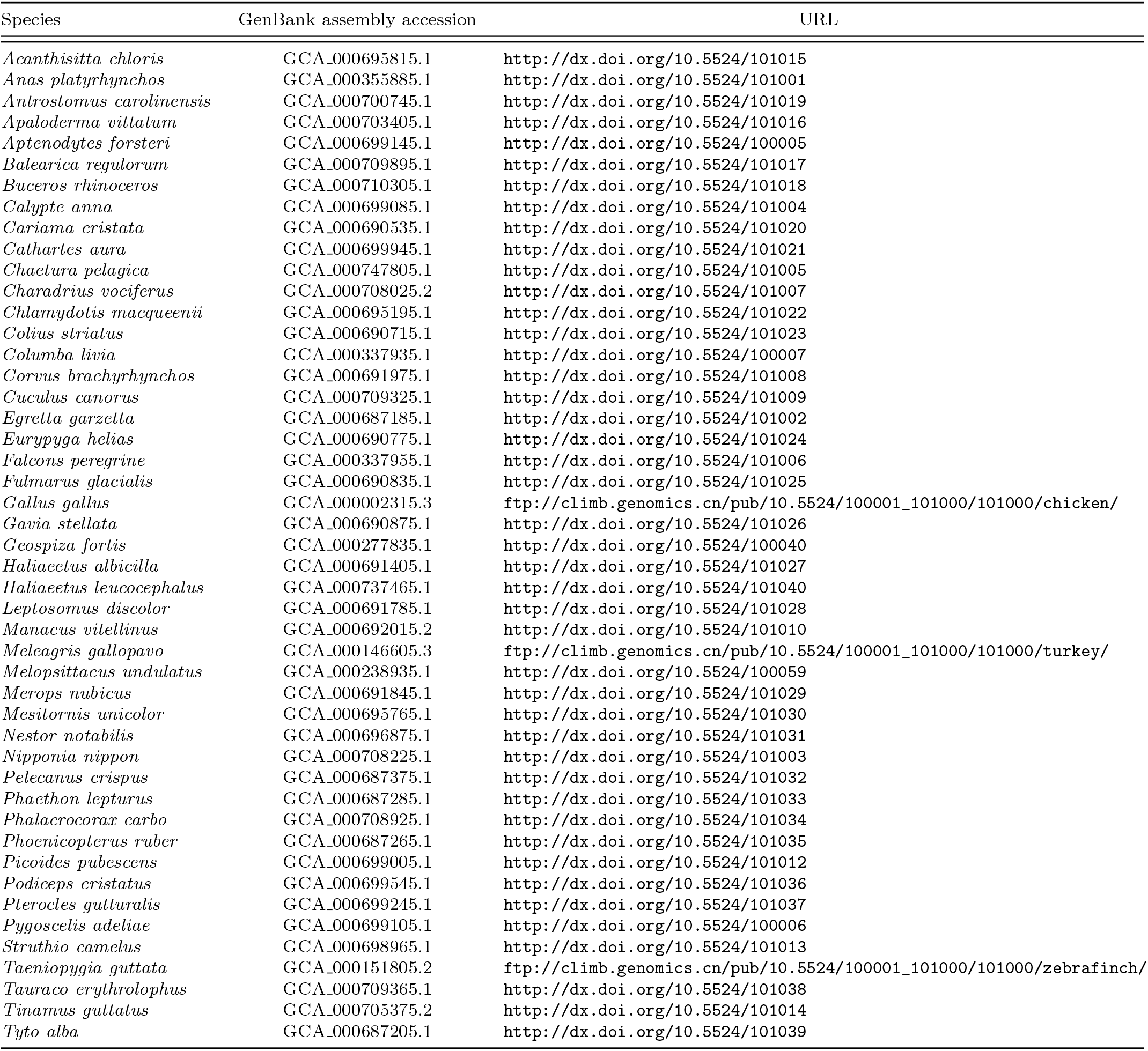
GenBank accession numbers and URLs for avian genomes

**Table S7:** The coverage of genomes over three datasets. Each genome is skimmed with 100Mb sequence.

| Dataset | Min | Mean | Max |
| --- | --- | --- | --- |
| Drosophila | 0.45X | 0.60X | 0.79X |
| Anopheles | 0.37X | 0.57X | 1.02X |
| Birds | 0.082X | 0.090X | 0.107X |

**Table S8:** Comparing the average error of Mash, Skmer, and AAF over three datasets. Fixed sequencing effort from each species.

| Dataset | Sequencing effort | Mash | Skmer | AAF (uncorrected) | AAF (corrected) |
| --- | --- | --- | --- | --- | --- |
| Anopheles | 0.1Gb | 48.02% (1.54%) | 2.02% (0.05%) | 40.22% (1.67%) | 9.62% (0.52%) |
|  | 0.5Gb | 24.89% (0.59%) | 0.75% (0.02%) | 17.60% (0.70%) | 7.35% (0.26%) |
|  | 1Gb | 18.43% (0.54%) | 0.55% (0.02%) | 16.94% (0.61%) | 4.74% (0.22%) |
| Drosophila | 0.1Gb | 47.98% (0.82%) | 1.65% (0.06%) | 40.67% (0.94%) | 9.00% (0.20%) |
|  | 0.5Gb | 25.25% (0.34%) | 0.72% (0.03%) | 18.63% (0.45%) | 7.00% (0.19%) |
|  | 1Gb | 13.00% (0.16%) | 0.50% (0.02%) | 19.69% (0.52%) | 2.18% (0.06%) |
| Birds | 0.1Gb | 95.57% (2.54%) | 5.72% (0.06%) | 86.45% (3.18%) | 49.48% (1.94%) |
|  | 0.5Gb | 56.61% (1.40%) | 2.14% (0.02%) | 49.13% (1.75%) | 13.73% (0.56%) |
|  | 1Gb | 41.25% (0.97%) | 1.32% (0.01%) | 34.33% (1.22%) | 1.05% (0.09%) |

\* The standard error of the mean is provided in parentheses.

